# Postsynaptic protein assembly in three- and two-dimensions studied by mesoscopic simulations

**DOI:** 10.1101/2023.03.09.531849

**Authors:** Risa Yamada, Shoji Takada

**Affiliations:** Department of Biophysics, Graduate School of Science, Kyoto University, Kyoto, Japan

## Abstract

Recently, cellular biomolecular condensates formed via phase separation have received considerable attention. While they can be formed either in cytosol (denoted as 3D) or beneath the membrane (2D), the underlying difference between the two has not been well clarified. To compare the phase behaviors in 3D and 2D, postsynaptic density (PSD) serves as a model system. PSD is a protein condensate located under the postsynaptic membrane that influences the localization of glutamate receptors and thus contributes to synaptic plasticity. Recent *in vitro* studies have revealed the formation of droplets of various soluble PSD proteins via liquid-liquid phase separation. However, it is unclear how these protein condensates are formed beneath the membrane and how they specifically affect the localization of glutamate receptors in the membrane. In this study, focusing on the mixture of a glutamate receptor complex, AMPAR-TARP, and a ubiquitous scaffolding protein, PSD-95, we constructed a mesoscopic model of protein-domain interactions in PSD and performed comparative molecular simulations. The results showed a sharp contrast in the phase behaviors of protein assemblies in 3D and those under the membrane (2D). A mixture of a soluble variant of the AMPAR-TARP complex and PSD-95 in the 3D system resulted in a phase-separated condensate, which was consistent with the experimental results. However, with identical domain interactions, AMPAR-TARP embedded in the membrane formed clusters with PSD-95, but did not form a stable separated phase. Thus, the cluster formation behaviors of PSD proteins in the 3D and 2D systems were distinct. The current study suggests that, more generally, stable phase separation can be more difficult to achieve in and beneath the membrane than in 3D systems.

**SIGNIFICANCE:** Synaptic plasticity is a key factor in memory and learning. Upon learning, protein condensates that form beneath the postsynaptic membrane are known to change their nature. Recent studies have suggested that condensate formation is related to liquid-liquid phase separation based on *in vitro* experiments of soluble parts. However, the phase behavior can be strongly dependent on physical dimensions. The mechanism by which condensate grows beneath the membrane is not well characterized. Taking advantage of the ease of systematic comparison using computer simulations, we investigated the phase behaviors of postsynaptic protein assemblies in 3D and 2D systems. The results revealed that even when a 3D system exhibited clear phase separation, the corresponding 2D system did not exhibit it stably.

## INTRODUCTION

Recently, biomolecular condensates in cells have received considerable attention (1). Biomolecular condensates are heterogeneous assemblies that are composed of many proteins and RNAs. As these condensates are considered to have cellular functions but are not surrounded by membranes, they are often called membrane-less organelles. Classic examples include Cajal bodies, nuclear speckles, stress granules, and postsynaptic density (PSD), whereas recent studies have revealed many more examples, such as signalosomes (2). These condensates are formed mainly by two types of interactions: interactions mediated by disordered regions with less sequence specificity, and specific and stoichiometric interactions between protein domains (3–7). Notably, these condensates can form either in solutions, as in the case of Cajal bodies (the 3D system), or under the membrane, as in the case of PSD (5, 8) (the 2D system). Biomolecular condensates often form via liquid-liquid phase separation (LLPS). It is well-known in statistical physics that phase transition behaviors are strongly dependent on the underlying physical dimension (9); 1D systems cannot show any phase separation, 2D systems are marginal, and 3D systems tend to exhibit sharp phase transitions. This trend may sound odd because LLPS in biology has often been reported in 2D systems. While 3D and 2D LLPS behaviors can qualitatively be different, the possible differences have not been well explored, to the best of our knowledge.

To compare the phase behaviors in 3D and 2D, PSD proteins can serve as a model system. PSD is a protein condensate beneath the postsynaptic membrane that was first observed by electron microscopy at the distal tip of the spine head (10). In hippocampal CA1 pyramidal cells, PSD has an area of 0.05–0.3 μm^2^ and a thickness of approximately 20 nm (11) containing several hundreds to thousands of proteins, as estimated from proteomic analysis (12–15). Proteins identified as components of PSD are diverse and have a hierarchical structure ranging from membrane-embedded proteins such as AMPA receptors (AMPARs) and cell junction proteins to cytoplasmic proteins, including scaffolding, signaling, and cytoskeletal proteins (15–17). As these proteins have diverse signaling pathways (18), PSD is regarded as a protein condensate specializing in signal transmission and synaptic plasticity. However, PSD protein condensates or clusters have been suggested to be driven by LLPS of major scaffolding proteins in cytosol (the 3D system) (8).

The scaffolding protein PSD-95, a member of MAGUKs, is one of the most abundant proteins in PSD, with approximately 300 PSD-95 molecules estimated to exist in each PSD (19, 20). PSD-95 is localized near the postsynaptic membrane in the upper part of the PSD hierarchy, and as a scaffolding protein, contributes to the formation and maintenance of protein condensates beneath the membrane. PSD-95 contains three PDZ domains, each of which has relatively weak one-to-one specific interactions with the carboxyl tail regions (the PDZ binding motif (PBM)) of target proteins, such as Shank, SAPAP, and TARP(21). The former two proteins further bind to proteins such as Homer, which are eventually associated with cytoskeletal proteins at the bottom of the PSD hierarchy. Thus, PSD-95 is involved in the facilitation of PSD condensation.

Glutamate receptors in the postsynaptic membrane are also components of PSD at the top of the PSD hierarchy and are directly involved in postsynaptic signaling. AMPAR, a representative transmembrane glutamate receptor, forms a tetrameric structure and interacts with PSD-95 via the regulatory protein TARP (transmembrane AMPA receptor regulatory protein). Specifically, each AMPAR subunit stoichiometrically binds to one TARP at its transmembrane regions. The C-terminal tail of TARP with PBM binds to PSD-95 in the cytoplasm beneath the membrane. Observations of postsynaptic membranes by STORM and others have shown that AMPARs are compartmentalized into their “nanodomains” (22). Simulations (23–26) suggest that not only the number of AMPARs but also the localization of AMPARs strongly affects the amplitude of excitatory postsynaptic currents (27). The spatial arrangement of AMPARs plays a key role in the efficiency of the postsynaptic response, and is controlled by the formation of protein condensates with PSD-95 via TARP.

Recent *in vitro* experiments have demonstrated that PSD-95 forms a liquid condensate with the C-terminus of stargazin, one of the subtypes of TARP (28, 29), or with NR2Bc, the C-terminus of NMDAR (8, 29) in solution, primarily via protein-domain interactions. However, most of the previous *in vitro* experiments demonstrating phase separation used soluble variants in the 3D system (28), and thus can be different from the behavior that would occur in and beneath the postsynaptic membrane. The detailed molecular mechanisms by which PSD-95 regulates AMPAR localization on the membrane remain unresolved.

The purpose of this study was to compare the phase behavior and interaction network of the protein condensate containing AMPARs, TARPs, and PSD-95 under the postsynaptic membrane (2D system) with a system containing soluble variants of the same proteins in solution (3D system) via computer simulations of mesoscopic models. The soluble system does not directly reflect the 2D system embedded in the postsynaptic membrane. The 2D system should be more restricted than the 3D system to form a dense protein network in the condensate. However, clarifying the differences between 2D and 3D systems through experiments is inherently difficult. Thus, in this study, we took advantage of computational modeling.

In this study, we constructed a mesoscale model in which each protein domain was simplified as a single spherical particle and performed comparative molecular simulations of AMPARs, TARPs, and PSD-95s under the membrane (2D system) and their soluble counterparts in the 3D system. The proposed domain-resolution mesoscopic model makes comparative simulations of the LLPS dynamics tractable (30, 31). In this model, specific interactions between protein domains are treated as virtual chemical bonds formed by reactions, which ensure one-to-one stoichiometric binding within the mesoscopic representation. The key parameters in the model were calibrated using a 3D system based on *in vitro* experimental data. We analyzed the protein domain interaction networks formed in the 3D and 2D systems, and found significantly reduced network edges in the 2D system. While the 3D system exhibited clear phase separation, scaling analysis of the 2D system indicated that the mixture of AMPARs, TARPs, and PSD-95 beneath the membrane formed finite-sized clusters, but not the separated phase. In general, the results suggest that the phase separation behaviors are markedly different between the 3D and 2D systems. The droplet formation in a soluble 3D system does not directly prove phase separation in the corresponding 2D system.

## METHODS

### Mesoscopic model: Molecular representation

Based on previous experimental assays (28, 29), we simulated the mixture of a glutamate receptor AMPAR (or its soluble counterpart), a regulatory protein TARP, and a scaffolding protein, PSD-95, using a mesoscopic model (30, 32, 33). In the mesoscopic model, each globular domain was represented by a single spherical particle. The neighboring domains are connected by interactions derived from the Gaussian polymer model, which represents flexible linkers.

In this study, we considered two simulation systems: 1) a system containing soluble variants of AMPAR and TARP, together with PSD-95s. AMPAR and TARP were replaced with DsRed2 (PDB ID: 1ZGO, *N* = 936 residues) and the soluble C-terminal part of TARP (TARPc), respectively, based on the experiment (29) (denoted as the 3D system) (Fig.1A top); and 2) the system that contained AMPAR and TARP restrained to the membrane surface, in addition to palmitoylated PSD-95 (denoted as the 2D system) (Fig.1B top). Thus, in addition to the underlying geometry, the two systems have different molecular sizes and diffusion coefficients. However, given the mesoscopic nature of the model, these differences would not severely alter the results.

**Figure 1.**
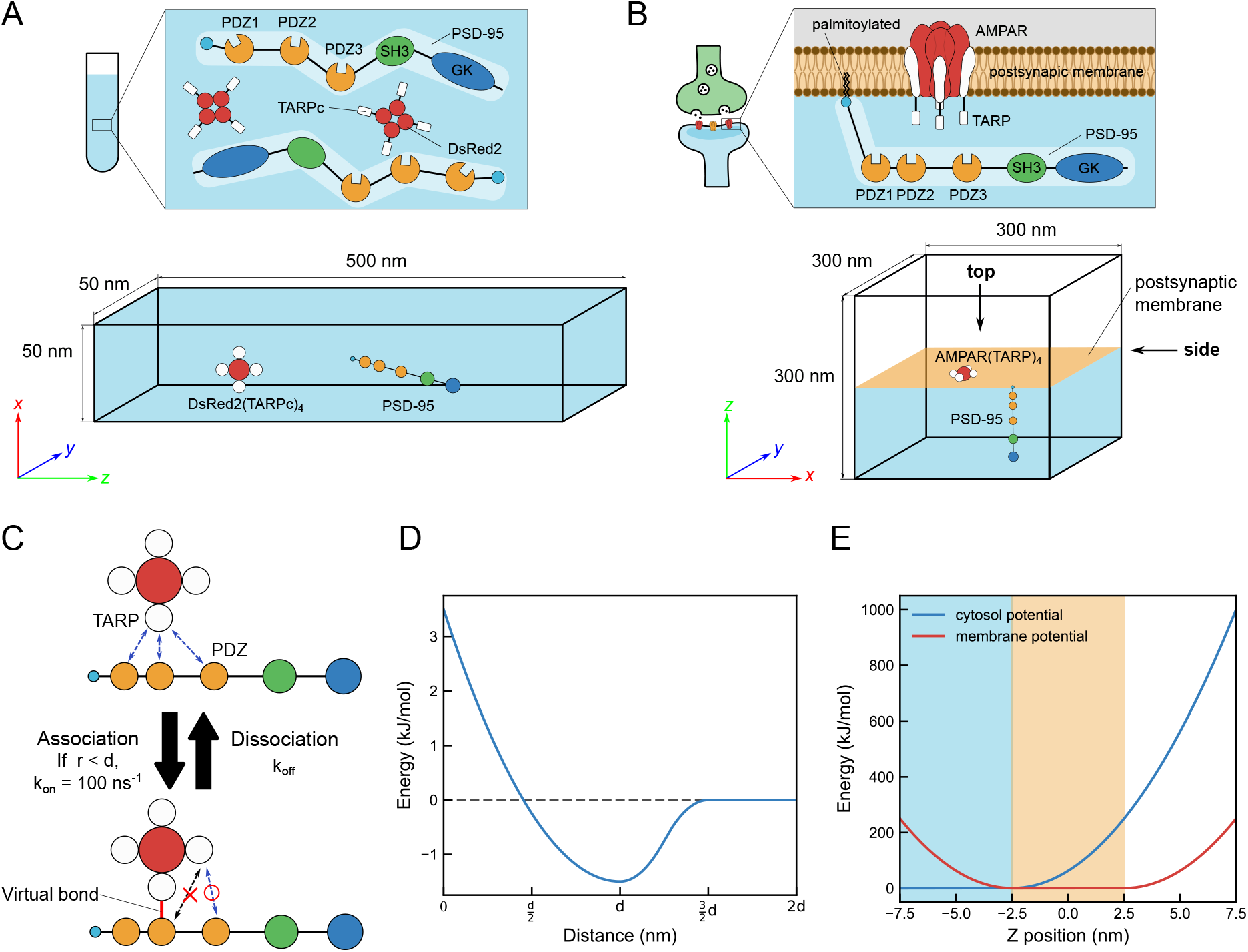
The mesoscopic model in 3D and 2D systems. A) Schematic of the 3D system (top) and the simulation box (bottom). Soluble DsRed2 tetramer and each C-terminus of TARP are modeled by one spherical particle. B) Schematic of the 2D system (top) and the simulation box (bottom). The AMPAR and each TARP are modeled by one spherical particle. PSD-95 is modeled by spherical particles with six connected particles: N-terminus, PDZ1, PDZ2, PDZ3, SH3, and GK domain. Particles representing AMPAR, TARP, and N-terminal of PSD-95 diffuse on the planar membrane (orange), while other particles diffuse in the cytosol just below the membrane (sky-blue). Among many, one representative box size is shown. C) The virtual reaction to represent one-to-one stoichiometric interaction (specific interaction) between TARP and PDZ domains. D) The non-specific inter-domain interactions. E) Implicit membrane potentials of the particles representing membrane embedded domains (red) and those representing cytosolic domains (blue). Sky-blue: cytosolic region, Orange: membrane region.

Although AMPAR forms a tetrameric structure constituted by GluA1-4 subunits, in this study, we regarded it as a single sphere, assuming that its tetrameric structure was maintained throughout the simulation. The AMPAR is assumed to be composed of a homo-tetramer of the GluA2 subunit (PDB ID: 3kg2, *N* = 3292 residues). TARP is a regulatory protein that binds to the GluA subunit on a one-to-one basis, thereby regulating AMPAR protein levels, synaptic localization, and channel activity (34). As a representative of TARP proteins, we chose the TARP γ2 subunit (*N* = 211 residues), also known as “stargazin” since it has been studied as the most well-known TARP subtype in numerous preceding researches. We modeled the TARP (or TARPc) protein as a single spherical particle. Throughout the simulation, we assumed that four TARP particles were always connected to one AMPAR (or DsRed2) and never dissociated from AMPAR. It can thus be denoted as DsRed2(TARPc)_4_ and AMPAR(TARP)_4_ for the 3D and 2D systems, respectively. PSD-95 contains five globular domains: three PDZ domains (PDZ1, PDZ2, and PDZ3, all *N* = 86 residues), one SH3 domain (*N* = 70 residues), and one GK domain (*N* = 165 residues) (28, 35). In addition, an additional small particle representing the first five amino acids of PSD-95 with two cysteine residues, which are palmitoylated in the case of 2D system, is added to the N-terminus (*N* = 5 residues) and tethered to the postsynaptic membrane (36). We represent a PSD-95 molecule as six spherical particles connected by a Gaussian chain. In the 3D system, a single sphere representing an AMPA receptor in the 2D system was substituted by a soluble fluorescent protein called DsRed2, which also has a tetrameric structure (29). The N-terminal particle of PSD-95 was also replaced with a sphere, with no restraint on the membrane.

The radii *R*_n_ (in nm) and diffusion coefficients *D* of globular domains were set using the formula below as a function of the number of amino acids *N* in the domain (37):

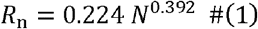

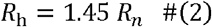

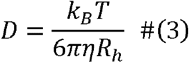

where *R*_h_ is the hydration radius, *K*_*B*_ is the Boltzmann constant, *K*_*T*_ is the temperature, and ηis viscosity. Since *R*_η_ is determined by the number of amino acid residues, molecular size of AMPAR in the 2D system is larger than that of DsRed2 in the 3D system for the representation of a tetrameric protein with 4 TARPs. Viscosity in the 3D system and that of the domains in the cytosol in the 2D system were determined by water viscosity at 300 K (0.89 cP). The viscosity of the membrane-embedded domains in the 2D system was set to be 100 times that of the cytosol, since the viscosity of the bilayer core in the DPPC vesicle was estimated to be 80-100 cP (38).

Mesoscopic model: Potentials

Based on the molecular representation in each system, the molecular system is subject to the total potential energy function as follows:

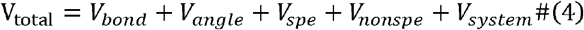

Where *V*_bond_ is of two types: harmonic bonds *V*_bond − h_ and interactions derived from the Gaussian polymer chain *V*_bond − g_. We set the bond between AMPAR and TARP as a harmonic bond. The spring coefficient of the harmonic bonds was set to 10 kJ/mol/nm^2^. On the other hand, protein linkers in PSD-95 are represented by a Gaussian polymer chain, which is easy to extend and shrink, reflecting the flexibility of the protein linker. Compared to that of the harmonic bond, which is set relatively high to maintain a stable tetrameric structure, the spring constant of the Gaussian chain was determined by the number of amino acids (*N*) contained in the linker and takes the value from k ∼ 0.3 to 3 in the current case.

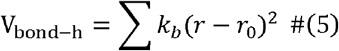

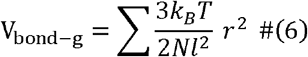

Where *k*_*b*_ is the spring constant of the harmonic bond (set as 10 kJ/mol/nm^2^), *r*_0_ is the natural length of the harmonic bond set as the sum of the radii of the interacting domains (corresponding to the sum of the AMPAR and TARP radii), *K*_B_ is the Boltzmann constant, *I* is the length between two amino acids (set as 0.38 nm), and *r* is the distance between neighboring domains. In addition, the angle potential *V*_angle_ was applied to TARP-AMPAR-TARP to reproduce the correct geometry of the complex.

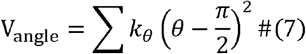

Where *kθ* is the spring constant (set as 10 kJ/mol/nm^2^ only to maintain the proper shape of AMPAR-TARP complex). It should be noted that the modeling only with the bond and angle potentials does not constrain the AMPAR(TARP)_4_ complex to the planar geometry, which might cause some artifacts. To test this possibility, we also implemented the model that additionally includes the dihedral angle potential constraining the planar geometry of the AMPAR(TARP)_4_ complex.

The *V*_spe_ represents the specific interaction between a TARP and PDZ domain. To accurately represent the one-to-one stoichiometric binding of the two domains, we used a virtual bond produced by a virtual reaction (Fig.1C). The virtual reaction is defined by the association and dissociation reactions. The association reaction occurs with a microscopic rate constant *K*_on_ when the distance between a TARP and a PDZ domain is within the threshold distance (set as *r*_0_, the sum of the TARP and PDZ radii). In the simulation, the association trial was examined every timestep Δ*t* (Δ*t* = 0.25 ns in the 3D system and 1.0 ns in the 2D system). In each trial, the association occurred with Poisson probability P = 1−exp(−*k*_on_ Δ*t*). In this model, the microscopic rate constant *K*_on_ was fixed at 100 /ns throughout all simulations. This rate constant value indicates that when two reactive molecules are in close proximity, a bond is created between them with a probability very close to unity. Thus, we effectively used the Smoluchowski model for diffusion-controlled reactions (39). The dissociation reaction occurred at a dissociation rate constant,*k*_off_. The equilibrium dissociation constant for the specific binding *k* _d,spe_ is not identical, but is correlated with *k*_off_/_on_. In this study, we enforced *k*_d,spe_ as the experimental value by altering *k*_off_, whereas *k*_on_ was fixed throughout. Without any specific knowledge on the binding/dissociation dynamics, it is reasonable to assume that the binding rate *k*_on_ is diffusion-controlled and thus is insensitive to the interaction strength, whereas the dissociation rate *k*_off_ depends on the intermolecular energy and is more directly linked to the equilibrium constant. When, and only when a TARP and a PDZ were bound (within the same “topology”), we turned on a harmonic interaction between the particles.

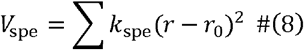

where *k*_spe_ was set to 10 kJ/mol/nm^2^, which is the same as that for other harmonic bonds. Here, we call the interaction defined as a virtual bond as a “specific interaction.” By appropriately defining virtual reaction systems, we can rigorously exclude the binding of PDZ domains to TARPs that are already bound to other PDZ domains, guaranteeing a one-to-one specific interaction. Although the C-terminus of TARP, which contains a PDZ binding motif, is said to interact mainly with PDZ1 or PDZ2 on the postsynaptic membrane(28), we assumed that all PDZ domains have the same binding ability to TARPc in this model, as the binding ability to PDZ3 has also been confirmed by artificially extending the C-terminal tail of TARP (35).

In addition to the virtual bond created by the virtual reaction, we implemented the nonspecific interaction *V*_nonspe_ which acts between any two protein domains, in contrast to the specific interaction. This interaction consists of a weak inter-domain force and an excluded volume term between the two particles. The potential function is obtained by connecting the three harmonic functions as follows (Fig.1D) (32):

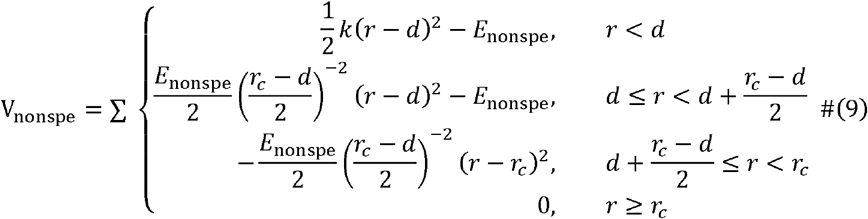

where the distance of the potential minimum *d* was set as the sum of the radii *r*_0_. The cutoff was set to 1.5 *r*_0_. The spring constant as set to 10 kJ/mol/nm^2^. The depth of the potential minimum,*E*_*nonspe*_ was tuned based on the experimental data, as described below. Note distance that even if *E*_*nonspe*_ is set to zero, a repulsive potential is applied when the intermolecular distance is smaller than and plays the role of excluded volume.

To reproduce and compare the situation in an *in vitro* experiment of soluble counterparts and the state of the mixture of AMPARs and PSD-95 in the actual postsynaptic membrane, we performed simulations in 3D and 2D systems.

In the 2D system, transmembrane domains of proteins (AMPAR, TARP, and the palmitoylated N-terminus of PSD-95) diffused within the membrane space (defined as | Z | < D_*mem2*_), while other domains diffused in the cytoplasm under the membrane (Z<− D_*mem2*_), where the half membrane thickness D_*mem2*_ was set to 2.5 nm. To achieve this, we introduced an additional potential that implicitly represents the membrane. We used transmembrane domains (Fig.1E, right).

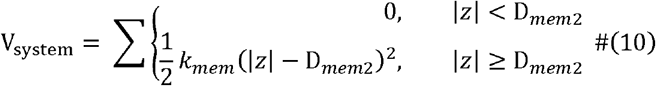

Where *K*_mem_ was setd to 10 kJ/mol/nm^2^. For cytoplasmic domains, we applied

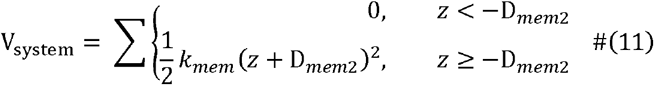

All potentials are made of soft-core potentials to realize efficient MD simulations for a dense mixture of proteins during liquid-liquid phase separation using a relatively large time scale. The results can depend on the value of *k*_*mem*_ to some extent. However, given the resolution of the mesoscopic model, we do not argue much on this subtlety. While TARP is modeled as a single sphere, TARP is tethered to the membrane via its transmembrane helix and the C-terminal tail of TARP interacts with the PDZ domain in the cytosol beneath the membrane.

The simulations were performed using ReaDDy 2 (32, 40). Protein dynamics were modeled using the overdamped Langevin equation with the temperature 300 K,

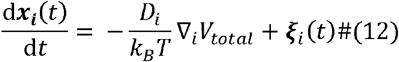

Where ***x***_*i*_ (*t*) ∈ ℝ^3^is the coordinate vector of *i*-th particle at time, and ξ*i* (*t*)is a Gaussian random vector with

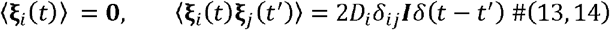

The time step of the simulation was 1.0 ns for the 2D system and 0.25 ns for the 3D system and the parameter tuning simulation. With time steps larger than these, we occasionally observed breakage of bonds between AMPAR and TARP. Given the simple Langevin dynamics used, the time scale in the software does not necessarily correspond precisely to actual physical time.

### Simulation Protocol: Slab simulation in the 3D system

To confirm whether the mixture of DsRed2(TARPc)_4_ complex and PSD-95 causes liquid-liquid phase separation through protein-domain interactions, we performed simulations in the 3D system and ran the simulation for 2 × 10^8^ MD time steps after 200 molecules of DsRed2(TARPc)_4_ and 200 molecules of PSD-95 were initially placed uniformly in a 50 × 50 × 500 nm slab box with the periodic boundary condition (Fig.1A bottom). Simulations were first performed using only specific interactions. The same simulation protocol was used to simulate LLPS with both specific and non-specific interactions.

As described above, the specific interaction between TARPc and PDZ is represented by a virtual reaction such that close molecules are tied together by virtual bonds. Here, we call the groups of molecules linked by virtual bonds as “clusters.”

### Simulation Protocol: Parameter tuning in the 3D system

We tuned the parameters for the specific and nonspecific interactions (*V*_spe_ +_nonspe_) based on experimental data. In the specific interaction, we fixed *k*_*on*_ and the threshold distance and altered *k*_*off*_ to control the specific interaction strength,*G*_*spe*_ = − *k*_*B*_ *T* In *k*d_,spe_. In nonspecific interactions, we tuned the depth of the potential well,*E*_*nonspe*_. To determine the balance between *G* _spe_ and *E*_*nonsp*e_, we conducted two assays.

The first assay aimed to reproduce the experimentally measured dissociation constant *K*_*d*_ between TARPc and PSD-95. The dissociation constant *K*_*d*_ of TARPc’s and PSD-95 in experiments ranged between 0.63–2.09 μM (28, 41) (we used 1.0 μM as the representative experimental *k*_*d*_ in this study). We conducted simulations of a single pair of molecules: one molecule each of PSD-95 and TARPc were included in a 50 × 50 × 50 nm cubic box. Using molecular simulations with various values of *G*_*spe*_ and *E*_*nonspe*_, the model dissociation constant was estimated using the following formula:

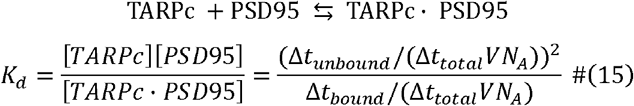

where Δ*t*_unbound_, Δ*t*_bound_, and Δ*t*_total_ are the cumulative durations of the unbound state, bound state, and total simulation time, respectively. *V* is the volume of the simulation box.*N*_A_ is Avogadro’s constant. In this tuning step, we assumed that two molecules are in “bound state” when the distance between TARPc and any PDZ domain is smaller than1.5 *r*_0_, which is the reference length at which two particles begin to interact. Conversely, “unbound state” is when the distances between TARPc and every PDZ domain are larger than the interaction length. This definition of bound/unbound state does not depend on the formation of a virtual bond created by specific interactions.

The second assay measured the concentration of the dilute phase when the system exhibited phase separation during the slab simulation. Experimentally, the critical concentration for phase separation, which corresponds to the concentration of the dilute phase in the phase-separated system, was ≾10μM. We note that DsRed2(TARPc)_4_ and PSD-95 have four and three interacting domains that interact one-to-one, respectively, and that the simulation box contains 200 molecules of each. This indicates that there is an excess of DsRed2(TARPc)_4_ as a client for specific interactions in this system. Therefore, we used the concentration of PSD-95 in the dilute phase as a reference for the dilute phase concentration. Adopting the slab simulation protocol described above, we repeated the molecular simulations with different parameter sets that provided appropriate *k*_d_ values. When well-separated phases were observed, we estimated the concentration of the dilute phase by counting the number of DsRed2(TARPc)_4_ and PSD-95 molecules in the dilute phase and averaging them with the final 2000 snapshots at each parameter set. Based on these values, we determined the values of *K*_off_ and *E*_nonspe_, which were compatible with the experimental *K*_d_ value and the critical concentration for phase separation.

### Simulation Protocol: Fluidity analysis in the 3D system

Two types of validation analyses, FRAP experiment-like analysis and mean square displacement (MSD) measurements, were performed to assess the molecular fluidity in the 3D simulation of the mixture of DsRed2(TARPc)_4_ s and PSD-95s.

In the FRAP experiment-like analysis, we first chose a reference time when most of the DsRed2(TARPc)_4_s and PSD-95s were involved in one large cluster. At the reference time, we marked the DsRed2 molecules in the left quarter (i.e., the negative z side) of the condensate. We tracked the marked molecules at subsequent times. The difference in the z-coordinates of the center of mass of the marked molecules and that of the rest was calculated as a function of time from the reference time point. The decay of the difference corresponds to the recovery of fluorescence in the FRAP experiment. We took an average of 1000 snapshots over the reference time point to obtain a smooth recovery curve.

In the MSD calculation, we first separated the slab box into condensed and dilute phases as described above. Then, in each phase, we obtained the MSD of the molecules as a function of time. Even though we used the periodic boundary condition, we could obtain the absolute distance of diffusion by tracking the motion. As in the FRAP experiment, we took an average of 1000 snapshots over the reference time point to obtain smooth MSD lines. The diffusion coefficients of the molecule in each phase were estimated using the value calculated from the slope of the MSD.

### Simulation Protocol: Simulation in the 2D system

We conducted a set of molecular simulations in and beneath the membrane of a mixture of AMPAR (TARP)_4_ and palmitoylated PSD-95. At the central plane of the 300 × 300 × 300 nm cubic box (the periodic boundary condition), we assumed an implicit membrane at z ∼0 and applied the implicit membrane potential to the particles to be embedded, as described above (Fig.1B bottom). Fifty AMPAR(TARP)_4_ complexes were uniformly placed on the membrane (z ∼0). Fifty PSD-95 molecules were uniformly placed beneath the membrane. The N-terminal beads of PSD-95 were in the membrane, while the other particles were in the cytosol (Z< −D). Starting from the configuration, we ran the simulation for 2 × 10^8^ MD time-steps. The same simulation was repeated thrice for statistical analysis. Using the same simulation protocol, we also executed simulations at three different areal densities (25 AMPARs/S, where S = 90000, 45000, and 22500 nm^2^) with the same number of AMPARs and PSD-95s. To monitor the stability of the clusters/condensates, we estimated the average number of molecules in the largest cluster for each configuration in the last 5,000 saved snapshots.

Next, with the areal density fixed to the dense value (25 molecules / 22500 nm^2^), we performed simulations containing different numbers of AMPAR(TARP)_4_ s and PSD-95s (n = 25, 50, 100, and 200). By investigating the scaling of the largest cluster size, we addressed whether the molecular assembly could be assigned as a separate phase. To assess the convergence of the cluster formation, we adopted a pair of simulations starting from mutually opposite extremes: one from a completely homogeneous configuration (denoted as a forward simulation) and the other from a completely phase-separated configuration (a reverse simulation. See Fig.S2 for the preparation of the initial state). We regarded the merging of the largest cluster size in the pair of simulations as a signal of equilibration. Beyond the first crossing time of the two-simulation data, we calculated the largest cluster size by averaging 100 consecutive snapshots at 50 independent time points.

## RESULTS AND DISCUSSIONS

### Model validation and phase separation in the 3D system

We first examined our mesoscopic modeling for the mixture of PSD-95 and DsRed2(TARPc)_4_, a soluble counterpart of AMPARs bound to the four C-terminal cytoplasmic parts of TARPs, based on available experimental data. This system has recently been shown to exhibit LLPS in *in vitro* assays. The C-terminus of stargazin (TARP _γ_2 subunit) and PSD-95 exhibit phase separation at an average concentration of 10 μM (28). In addition, the dissociation constant *K*_*d*_ between C-terminus of stargazin and full-length PDZ-95 was measured to be 0.63 ± 0.6 _μ_M (28), 1.2 _μ_M (result from ITC assay), or 2.66 ± 0.07 _μ_M (result from FP saturation assay) (41). We assumed the experimental *k*_*d*_ to be 1.0 _μ_M. These experimental data can be used for calibration.

We adopted the slab simulation box, which is known to be advantageous for realizing phase separation in MD simulations (42, 43). With more standard cubic boxes, protein condensates would take near-spherical droplets which are entirely surrounded by interface. The free energy cost from the interface results in large finite-size artifacts. The slab shape with the periodic boundary condition enables the separation of the two phases by the interfaces of small areas, which alleviates the finite-size effects (42, 43). We began the simulations using homogeneous and random configurations.

We first tested modeling solely by the specific one-to-one interaction between each of the PDZ domains of PSD-95 and TARPc. We set the nonspecific interaction to zero, i.e., *E*_nonspe_ =0, and tuned the strength of the specific interaction *G*_*spe*_ so that the computed dissociation constant k_d_ in the system of a single pair of molecules agreed with that of the *in vitro* experiment (see the Methods section for details). Using the calibrated *G*_spe_ (= 34.5 kJ/mol,*K* _off_ = 10^−4^ /ns) in the specific-interaction-only model, we performed a slab simulation containing 200 molecules of PSD-95 and DsRed2(TARPc)_4_ at a concentration of 267 μM, which is much higher than the experimental critical concentration of ∼10 μM. The simulation showed that the mixture did not exhibit phase separation, as illustrated in the top panel of Fig.2A. Thus, this specific interaction-only model contradicts experimental results.

**Figure 2.**
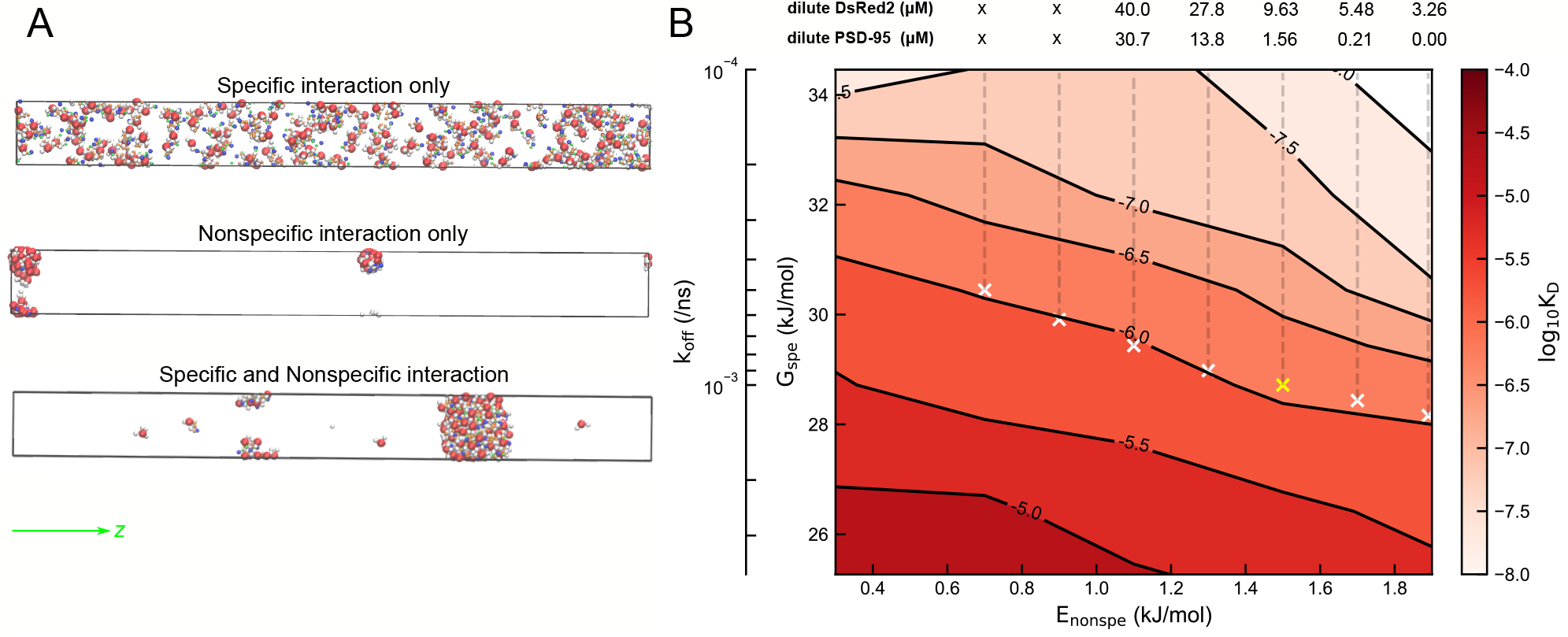
Model validation and the phase separation of PSD-95 and DsRed2(TARPc)_4_ complexes in the 3D system. A) Final snapshots in the three cases. (top) The case that contains the specific interactions alone. (middle) The case that contains nonspecific interactions alone. (bottom) The case that contains the balanced specific and nonspecific interactions. B) The computed _d_ of PSD-95 and DsRed2(TARPc)_4_ in the parameter space (*E*_*non* spe_ *G*_spe_) (contours and red color) and the concentration of the dilute phase in the phase separated slab system on the counter of the computed phase separation). The yellow cross mark indicates(*E*_*non* spe_ *G*_spe_) =(1.5, 28.7) (kJ/mol), the that agrees with the experiment (numbers above the figure, w here”x” m ea n the absence of the parameter we adopted. The leftmost axis is that of the *k*_off_ corresponding to *G* _spe_.

In the opposite limit, we removed the specific interaction and tuned the strength of the nonspecific interaction *E*_nonspe_ so that it agreed with the experimental *K*_d_ in the limit of the dilute condition. Using the calibrated *E*_nonspe_ (= 6.5 kJ/mol), we performed the same simulation as mentioned above, except for the interaction parameter. We found that PSD-95 and DsRed2(TARPc)_4_ formed small clusters, which gradually coalesced into larger clusters (a final snapshot in the second panel of Fig.2A). At the end of the 20 ms (8 × 10^7^ MD time step) simulations, we found no molecules in the dilute phase. We note that the concentration of the dilute phase corresponds to the critical concentration for phase separation (44), which indicates that the critical concentration of LLPS in this setup is much lower than 1 μM. This is inconsistent with the experimental results.

We then explored the two-dimensional parameter space (*E*_nonspe,_ *G*spe). First, we found a curve in which the computed *K* d agreed with the experimental value (a contour with white crosses in Fig.2B). Along the curve, we repeatedly conducted simulations of the mixture in the slab box from a completely phase-separated configuration and estimated the concentration of the dilute phase when the system maintained the phase separated state. The concentration of the dilute phase was estimated first by classifying bins along the long slab axis into dilute and condensed phases and then by determining the volume and number of molecules in the region classified as the dilute phase (The convergence of the simulation was examined in Fig. S1A). Along the curve on which the computed *k*_*d*_ agrees with the experimental one, at *E*_*n*onspe_ < 0.9 kJ/mol, the system did not maintain the phase separation resulting in uniform distribution (Fig.2B). On the other hand, with *E* _*nonspe*_ ≥0.9 kJ/mol, we found clear phase separation and a decrease in the concentration of the dilute phase as E_*nonspe*_ increased. Given tha t the critical concentration should be less than 10 μM, we chose (*E*_*n*onspe,_ *G* _nonspe_)=(1.5,28.7) (kJ/mol) as the optimal set of parameters (a final snapshot in the bottom panel of Fig.2A). Here, *k*_off_ is set to 10^−3^ /ns.

These results clarified that both the dissociation constant *K*_d_ in the dilute phase and the critical concentration of the phase separation can be simultaneously matched with the experimental data only when both specific and nonspecific interactions are implemented for the mixture of PSD-95 and DsRed2(TARPc)_4_. This finding is consistent with the results of recent studies (45, 46). Well-balanced combinations of specific and nonspecific interactions are necessary to describe phase separation in mesoscopic modeling.

Figure 3 shows the phase separation dynamics of a mixture of 200 PSD-95 and 200 DsRed2(TARPc)_4_ simulated in the slab box with the appropriately tuned parameter set. From a homogeneous and random configuration, many small and nearly spherical clusters of PSD-95 and DsRed2(TARPc)_4_ appeared in < 10 μs, which coalesced into larger and larger droplets, and finally became one large condensed phase together with the low but non-zero density phase at 40 ms (three snapshots in Fig.3A and movie in the supplemental data, Movie S1). Figure 3B shows the time course of the number of virtual bonds, i.e., the one-to-one binding between TARPc and PDZ (blue) and the number of DsRed2 (red) in the largest cluster. Since the time course of the cluster size has reached a plateau, we considered that the simulation reached the equilibrium at ∼40 ms. The increase in virtual bonds corresponds to the progress of domain-wise protein network formation, whereas the largest cluster size represents the progress of condensed phase formation. The state of the virtual bonds converged to equilibrium much faster than the phase behavior. As the fusion of two droplets doubles the number of particles in the largest cluster, we observed that the largest cluster grew step-by-step during the simulation. The coefficient of variation of the largest cluster size (Fig.S1B) shows that once a droplet is formed, it maintains the composition stably with little fluctuation.

**Figure 3.**
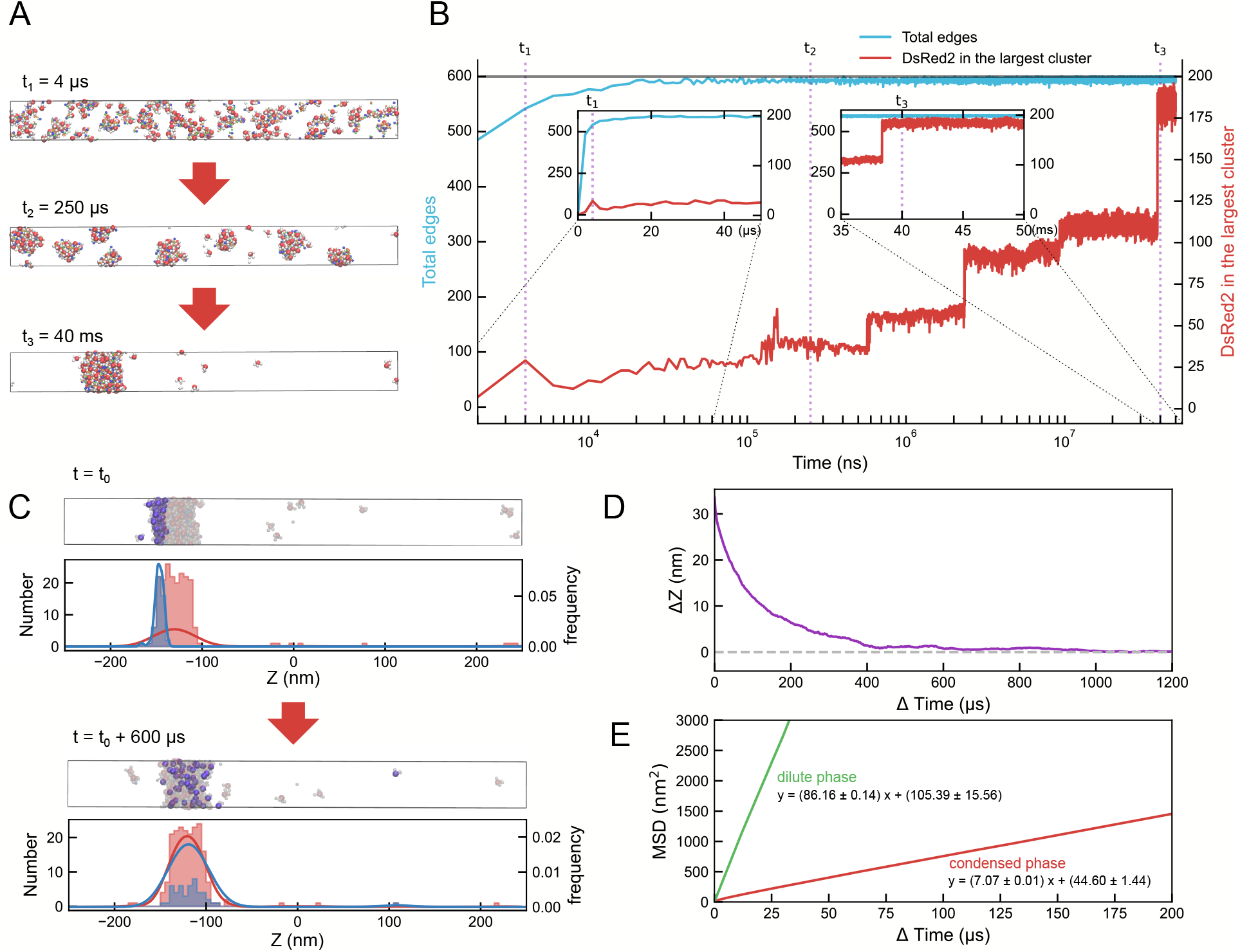
Phase separation dynamics for the mixture of 200 PSD-95 and DsRed2(TARPc)_4_ complexes in the 3D system. A) Snapshots at the three time points. B) Time series of the number of domain-wise specific bindings, virtual bonds (blue) and the number of DsRed2 in the largest cluster (red). The vertical dashed lines represent the time points at which snapshots are depicted in A. C) The FRAP-like analysis. The distribution of marked and the remaining DsRed2s at the reference time (top) and at 600 μs after the reference (bottom). D) Averaged decay of the difference between the center of mass of marked (photobleached) DsRed2s and that of the other DsRed2 particles. E) The MSDs of DsRed2 in the condensed phase (red) and in the dilute phase (green), respectively, as a function of time difference.

Notably, after the coalescence of two spherical droplets at early stages, the double-sized droplet rapidly assumed a nearly spherical shape. This is a hallmark of droplet fluidity.

To further assess whether the formed condensate has fluidity, we conducted fluorescence recovery after photobleaching (FRAP)-like analysis and calculated the MSD. FRAP experiments are often used to address the diffusivity of specimen molecules (47, 48). Recent FRAP experiments for mixtures of TARPc and PSD-95 showed that light photobleached fluorescence was restored, indicating the fluidity of the droplet (28). By mimicking the experiment, we marked DsRed2 molecules in a quarter volume of the condensed phase localized at one end (along the z-axis) at a reference time _O_ (purple at the top of Fig.3C) and tracked their movements in a subsequent time period. The DsRed2s molecules spread to the entire droplet after 1.2 ms (bottom of Fig.3C). To quantify the relaxation dynamics, we monitored the difference along the z-axis between the geometric centers of the marked DsRed2 and the other DsRed2 as a function of time from the reference time (Fig.3D), and found an exponential decay. This indicates that the formed condensates were liquid-like. We further calculated the MSD of DsRed2(TARPc)_4_ particles inside and outside the condensate (Fig.3E). The MSD curves of molecules belonging to either phase showed a linear dependence with respect to the MD time step. This means that molecules belonging to the two phases show normal diffusion with distinct diffusion coefficients. This is a hallmark of liquid droplets (49). Fitting MSD curve by linear regression, we estimated the slope of each MSD curve as 86.16±0.14 nm^2^/μs and 7.07±0.01 nm^2^/μs for the dilute and condensed phases, respectively. The diffusion coefficients of each phase were evaluated as 14.4±0.02 μm^2^/s in the dilute phase and 1.18±0.00 μm^2^/s in the condensed phase. An order-of-magnitude slower diffusion in the condensed phase is qualitatively consistent with the experiment though, quantitatively, the estimated diffusion coefficient of the condensed phase was about an order of magnitude larger than that can roughly be estimated from the FRAP experiment (28). This discrepancy could be attributed to somewhat arbitrary choices in the virtual reaction kinetic parameters; both protein binding and unbinding rates might be too large. However, this choice would not qualitatively change the behavior of a phase-separated condensate. Thus, we conclude that we have successfully built a mesoscopic model that reproduces the LLPS phenomenon with the soluble counterpart of the three-component postsynaptic density system (DsRed2-TARPc-PSD95) by combining specific interactions defined by virtual reactions and weak nonspecific interactions.

### Cluster formation on and beneath the membrane (the 2D system)

To address the difference between the assembly of the soluble counterpart of AMPAR, i.e., DsRed2, in *in vitro* experiment and that of AMPAR on the postsynaptic membrane, we performed simulations in a 2D system. In contrast to DsRed2(TARPc)_4_, AMPAR and TARP particles were restrained to the membrane. In addition, the N-terminal particle of PSD-95 was assumed to represent the palmitoylated group and was thus restrained to the membrane. These membrane-embedded particles have smaller diffusion coefficients in the membrane than their soluble counterparts in the solution. For interactions between TARP and the PDZ domain of PSD-95, we used the same parameter set as that used in the 3D system.

We first performed molecular simulations with 50 PSD-95 and 50 AMPAR(TARP)_4_ on the 300 × 300 nm membrane, which falls into a range of typical areal densities of AMPARs in the postsynaptic membrane (see below). The results showed that many small complexes of a few PSD-95s and AMPAR(TARP)_4_ appeared quickly (< 1 ms), which then coalesced into larger but unstable clusters (∼ 160 ms). A few proteins remained in the dilute phase (three snapshots in Fig.4A and movie in the supplemental data, Movie S2). From a random configuration, the number of domain-wise specific bindings (the network edges) increased rapidly (Fig.4B) over the timescale of 16 μs – 16 ms to reach a plateau. The number of AMPARs in the largest cluster grew at a later stage of ∼160 ms, as shown in the right snapshot of Fig.4A. Since this has reached a plateau, we judged that the simulation has reached the equilibrium at ∼160 ms. The largest cluster size converged to ∼20–30, which is approximately half the total number of AMPARs. In the same way as the 3D case, the one-to-one domain-wise binding of TARPs and PDZs progressed before the growth of the largest cluster size. Comparing the timescales, we found that domain-wise bindings grew more slowly in the 2D case than in the 3D case. This slower domain-wise binding can be attributed to smaller diffusion coefficients in the membrane. The increase in the largest cluster size (number of AMPARs) was also found to be stepwise. The time course of the coefficient of variation of the largest cluster size (Fig.S1C) shows that the droplets formed were easily broken up and repeatedly fused and split resulting in relatively large fluctuation. We plotted the distributions of the six domains of PSD-95 along the z-axis in Fig.4C, which shows that the domains closer to the N-terminus were, on average, located closer to the postsynaptic membrane. Although the spheres of the N-terminus of PSD-95 moving on the membrane due to palmitoylation can move freely in the membrane region (−2.5 nm < z < 2.5 nm), they are most abundant at z ∼ - 2.5 nm since N-termini are likely to be pulled down by other PSD-95 domains. Spheres representing PSD-95 domains other than N-termini were distributed in the range -15.0 nm < z < -2.5 nm below the membrane region. Although the existing ranges of the three PDZ domains (PDZ1, PDZ2, and PDZ3) were highly overlapped, the peaks of their distributions moved slightly away from the membrane from PDZ1, which is closest to the membrane, to PDZ3, which is farthest from the membrane, suggesting that the distribution of molecules to the postsynaptic membrane correlates with their distance from the protein N-terminus.

**Figure 4.**
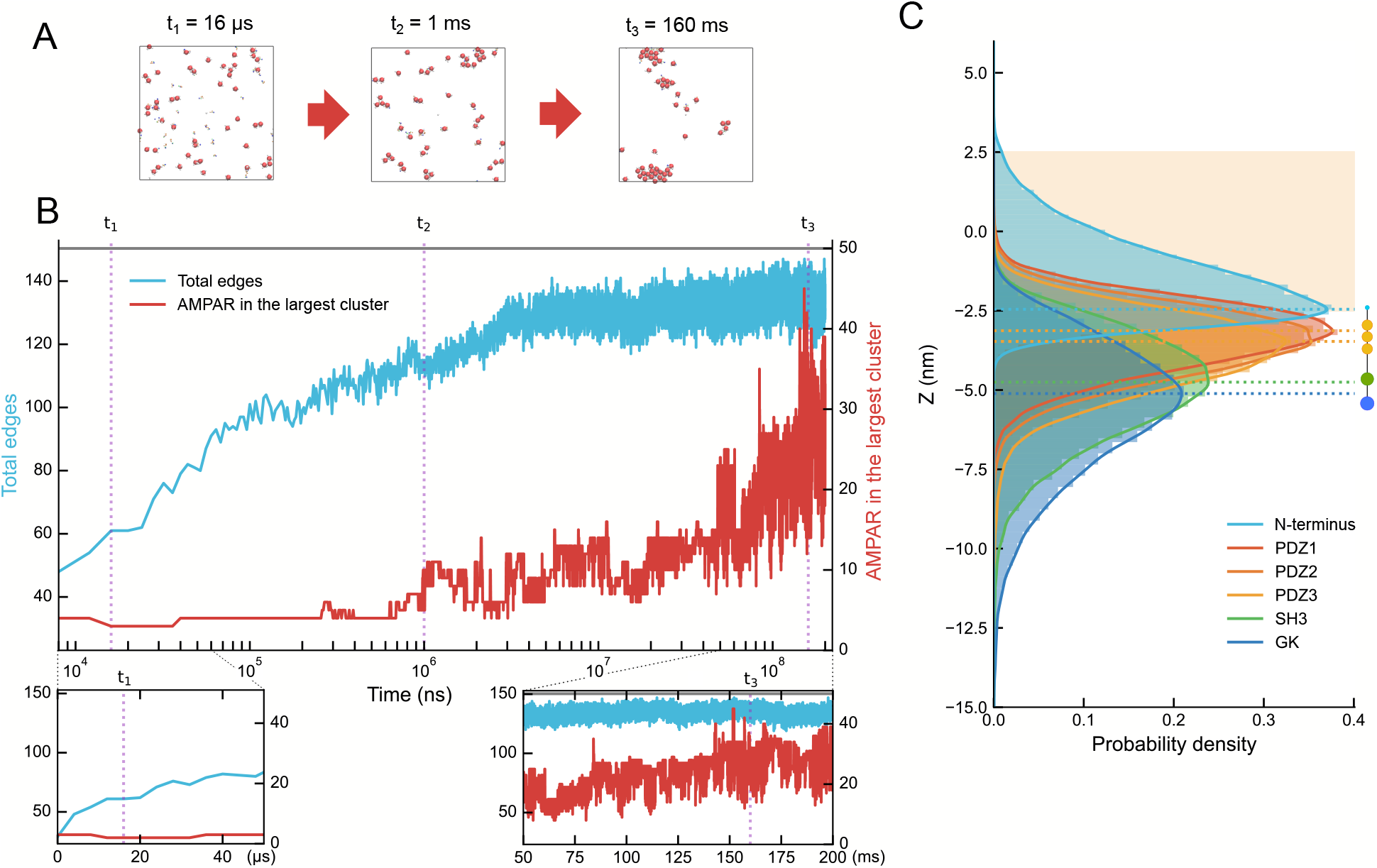
Molecular assembly dynamics for the mixture of 50 palmitoylated PSD-95 and 50 AMPAR(TARP)_4_ on and under the membrane of 90000 nm^2^ area (the 2D system). A) Snapshots at the three time points (t= t_1_, t_2_, t_3_). B) Time series of the number of domain-wise specific bindings, virtual bonds (blue) and the number of AMPARs in the largest cluster (red). The vertical dashed lines represent the time points at which snapshots are depicted in A. C) The distribution of domains in PSD-95 beneath membrane. The region -2.5 nm < z < 2.5 nm corresponds to the membrane (orange).

Next, we performed the simulations with three different areal densities: sparse condition (25 AMPARs/90000 nm^2^), medium condition (25 AMPARs/45000nm^2^), and dense condition: (25 AMPARs/22500nm^2^). The areal densities of AMPARs in the sparse and medium conditions correspond to typical areal densities at the postsynaptic spines of the adult rat hippocampus (50) and adult rat cerebellum (51, 52), respectively. A dense condition was used to investigate the limiting case.

Figure 5A shows the final snapshots of the simulation from a random and homogeneous configuration in the three densities from two directions, viewpoints **top** and **side** in Fig.1B (bottom). The results showed that PSD-95s assembled AMPARs to form clusters on the membrane at any areal density. Viewpoint **side** shows that the formation of AMPAR clusters (red particles) was co-localized with the PSD-95 molecules beneath the clusters. The temporal changes in the largest AMPARs clusters under three different areal density conditions (Fig.5B) showed that the lower the density, the longer it takes for the molecules to assemble, mainly as the probability of encountering molecules decreases in the lower density condition. While all the curves seem to converge by ∼ 60 ms, the largest cluster size continues to fluctuate significantly. Thus, the clusters were marginally stable with repeated coalescence and division, regardless of the areal density. This is in sharp contrast to the 3D case where we did not observe the division of large condensates. Under dense conditions, the majority of AMPARs (on average, 19.0 out of 25 AMPARs, Fig.5C right) were incorporated into the largest cluster, which might imply phase separation. On the other hand, with the medium and sparse conditions, no such large single cluster was formed stably, and two clusters containing about half of the total number of AMPARs (12.6 out of 25 AMPARs in the sparse condition, 11.2 out of 25 AMPARs in the medium condition, Fig.5C, left and middle, respectively) were observed for most of the simulation time. Under all conditions, we observed that some AMPAR included in a cluster occasionally left the cluster, diffused freely, and were absorbed into another cluster. In addition to AMPARs belonging to large clusters, single or small groups of AMPARs were also observed on the membrane. The features of cluster formation in the 2D system, such as stepwise growth of the cluster and slower domain-wise binding, were shared with all three areal density cases.

**Figure 5.**
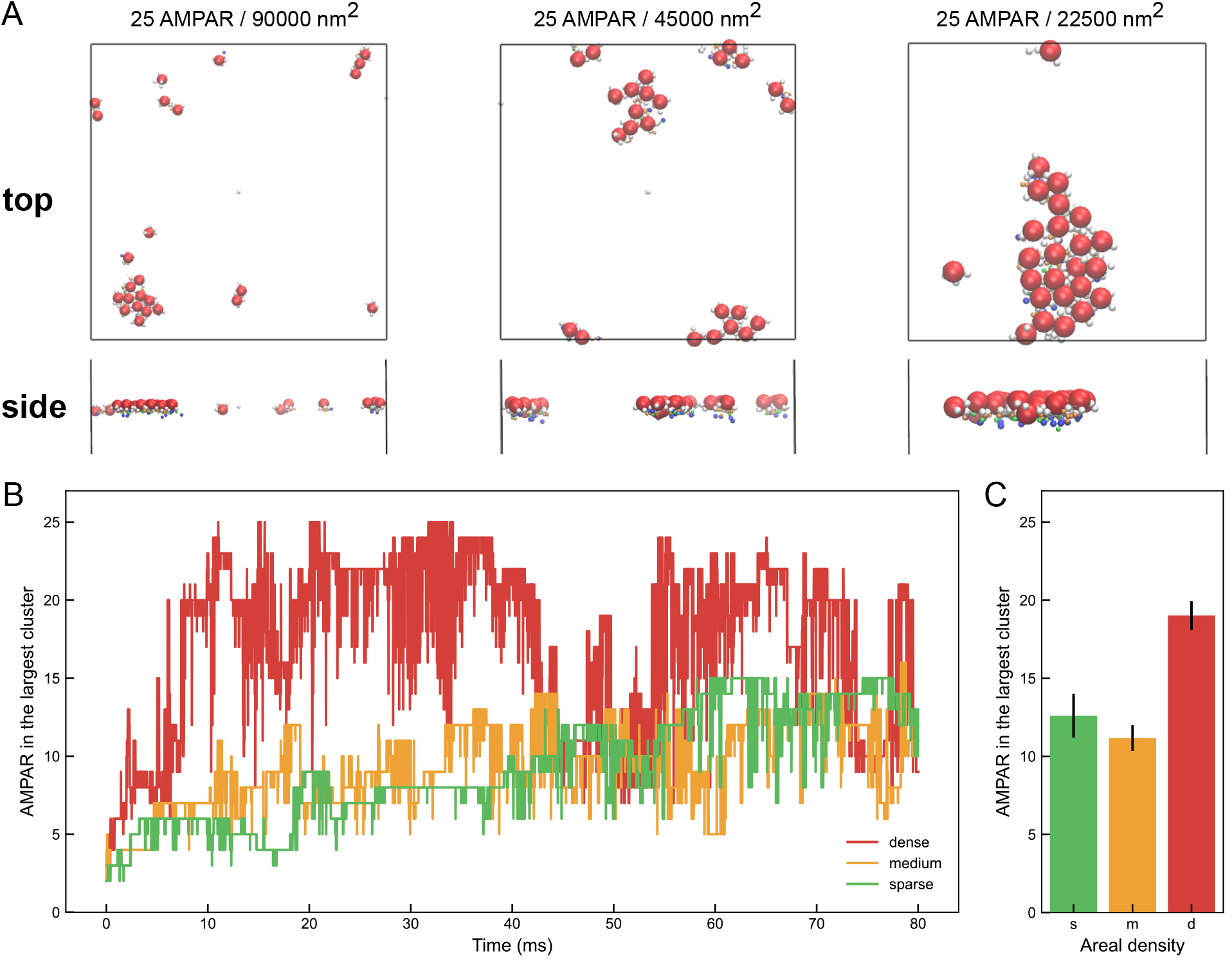
AMAPR(TARP)_4_ and palmitoylated PSD-95 assembly on and beneath the membrane in three different areal densities. A) Final snapshots of the protein assembly in three areal densities: the sparse (left), the medium (center), and the dense (right) conditions. Top and bottom pictures are views from the top and from the side, respectively. B) Time series of the number of AMPARs in the largest cluster for the case of the sparse (green), the medium (orange), and the dense conditions (red). C) The mean and the standard error of the largest AMPAR cluster size for the three areal densities.

### Receptor-scaffold network does not exhibit the phase separation on the membrane

Although both the mixture of DsRed2(TARPc)_4_ and PSD-95 in the 3D system and that of AMPAR(TARP)_4_ and PSD-95 in the 2D system formed clusters, we also observed that the stability of the large clusters differed between the two systems. Large clusters formed on and beneath the membrane were fragile, and breakage was often observed. On the other hand, we observed virtually no breakage of large clusters formed in the 3D system. In addition, the observed cluster sizes in the 2D system differed among the areal densities: a large cluster of receptors constituted by almost all the receptors in the system was observed in the dense condition but not in other conditions. While phase separation was clearly visible in the 3D system, it is unclear whether the cluster formation of receptors and scaffold proteins in the 2D system can be regarded as phase separation.

To address the phase behaviors more rigorously, we examined the scaling behavior both in the 3D and 2D systems. We note that for a cluster (droplet) formed in a multicomponent system to be regarded as a “phase,” the size of the phase should grow linearly with the system size in a set of self-similar systems (53, 54). Thus, we examined the self-similarity of droplets in a series of 3D and 2D systems of different sizes with identical densities (In the 2D system, we employed the dense condition). Both in 3D and 2D systems, we repeated simulations for the mixture of AMPAR-(TARP)_4_ and PSD-95 for four different numbers of AMPARs and PSD-95s at 25, 50, 100, and 200, and measured the number of AMPARs in the largest cluster. We note that, in the 3D system, while AMPAR is all substituted by DsRed2, we denoted it as AMPAR for simplicity.

To check the convergence of simulations, which was not easy particularly for larger systems, we employed a pair of simulations for each system size: a run from a random and homogeneous configuration (termed “forward”) and another run from a perfectly phase-separated configuration (termed “reverse”). Forward simulations are expected to show cluster growth, whereas reverse simulations can exhibit cluster breakage. The trajectories from the two starting configurations are expected to sample the identical ensemble when they reach equilibrium. For the fixed density, we show a pair of trajectories for each system containing 25, 50, 100, and 200 AMPAR(TARP)_4_s and PSD-95s (Fig.6A).

**Figure 6.**
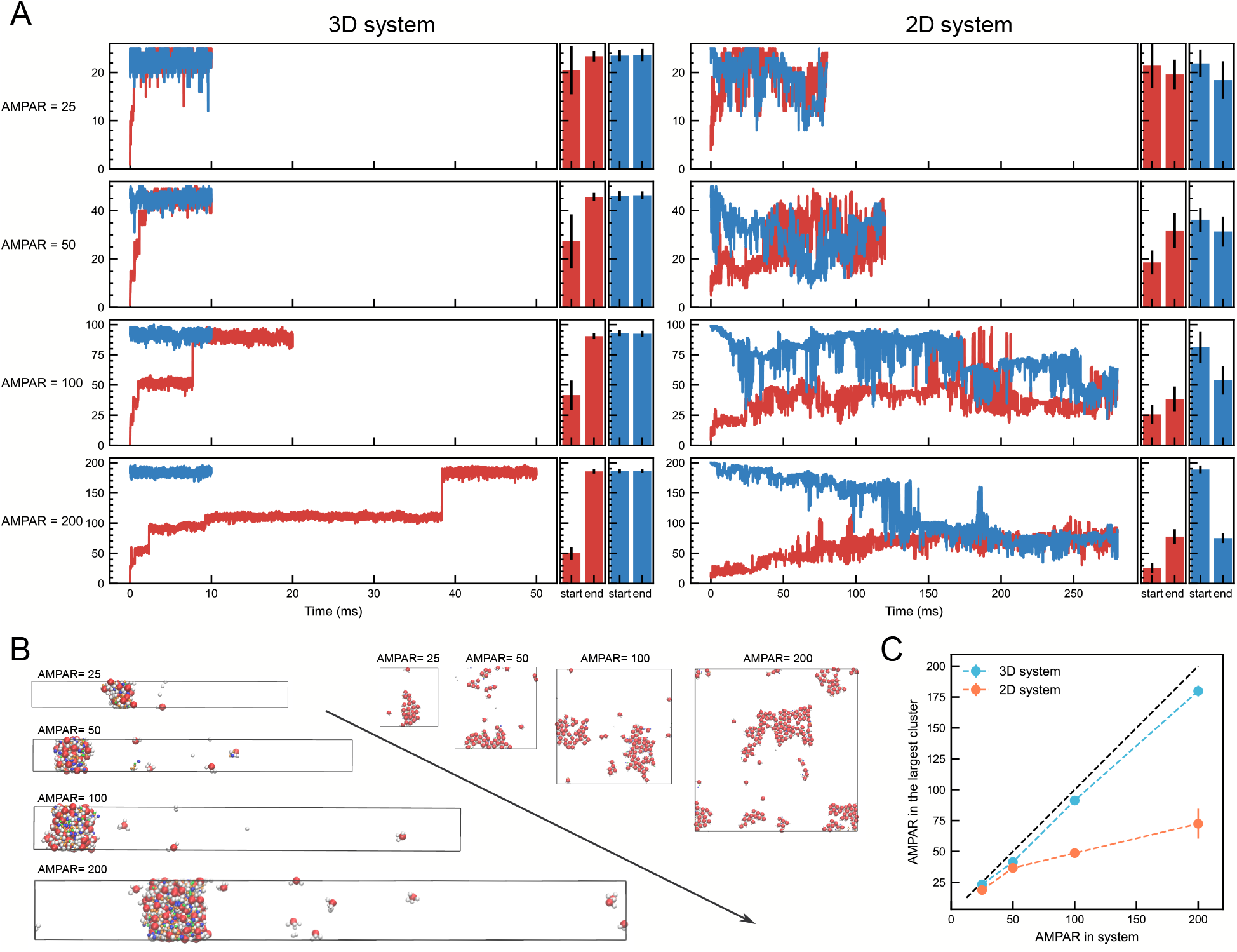
The scaling analysis of phase separation in the 3D and 2D systems for the fixed density (the dense condition in the 2D case). A) The forward (red) and reverse (blue) trajectories of the largest DsRed2/AMPAR cluster size as a function of time for four different system sizes of self-similarity; 25, 50, 100, and 200 AMPAR(TARP)_4_s and PSD-95s for the 3D (left) and 2D (right) systems (left panel of each system). The mean of the largest clusters at the beginning (denoted as start) and the end (denoted as end) of each simulation (right panel of each system). Bars; the standard error. B) The final snapshots of the four systems (left: 3D system, right: 2D system). C) The mean of the largest AMPAR cluster size as a function on the total number of AMPARs in the 3D (cyan) and 2D systems (coral). Black dashed line is the line assumed that the cluster growth is proportional to the cluster size in the system.

In the 3D system, the forward simulations show that the time course of the largest AMPAR cluster size reached plateau at approximately 1, 2, 8, and 40 ms for the systems with 25, 50, 100, and 200 AMPARs, respectively. We found the reverse simulations all reached to the largest cluster size equivalent to the plateau values in the forward simulations within 10 ms. In all the system sizes, the converged size of the largest cluster was only slightly smaller than total number of molecules in the system. Thus, majority of molecules were involved in the condensate (Fig.6B left).

In the 2D system, for a pair of forward and reverse simulations, we defined the time of convergence as the time when the largest cluster size in the forward simulation exceeds that of the reverse 50 time points. The average of three trials showed the time of convergence at ∼10, 40, 170, and 210 ms for 25, 50, 100, and 200 AMPARs, respectively. The distributions of the largest AMPAR cluster size after merging the two trajectories (the right panels in Fig.6A) and the evolutions of coefficient of variation (Fig.S2E) show broad distributions in the 2D system, indicating large intrinsic fluctuations. In the 2D system, snapshots in the final simulation step (Fig.6B right) suggest that the molecules form essentially one cluster for the case of 25 AMPARs but have several clusters for the case of 200 AMPARs. This is in sharp contrast to the geometrically similar growth of one large condensate in the 3D system.

Finally, we plotted the average size of the largest AMPAR cluster as a function of the number of AMPARs in the system (Fig.6C). The results indicate that although the largest AMPAR cluster size increases in both systems, the increase is not proportional to the number of AMPARs in the 2D system, contrary to the linear growth in the 3D system. The largest AMPAR cluster size in the 2D system seemed to saturate below ∼100. Sublinear growth and plausible saturation of the cluster size are hallmarks of the lack of phase separation. To assess the finite-size effect of the simulation box, we also performed a 2D simulation in a 2D-variant of the slab box finding that the scaling of the largest AMPAR cluster size is not significantly affected by the different box setup (Fig.S3). We also confirmed that there is little effect on the scaling of the largest AMPAR cluster size even under the planar constraints to the geometry of four TARPs, by introducing additional bonds and dihedral potentials in AMPAR(TARP)_4_ (Fig.S4).

We note that, in the 2D system, the areal density under this condition is probably larger than that of AMPARs on the actual postsynaptic membrane, as mentioned above. Even under such over-dense conditions, stable phase separation was not observed. With physiological AMPAR density, phase separation of the mixture of AMPAR(TARP)_4_s and PSD-95s is less likely to occur. Therefore, we suggest that any AMPAR density found in the postsynaptic membrane, such as in the hippocampus or cerebellum, is unlikely to cause stable phase separations on and beneath the membrane with only the binary mixture of AMPAR(TARP)_4_s and PSD-95s. Indeed, in an experiment in which receptor-like proteins were tethered on one side of the lipid bilayer to mimic PSD diffusion beneath the membrane, removal of even one of the four scaffolding proteins (PSD-95, GKAP, Shank3, Homer3) was found to completely prevent PSD cluster formation, supporting our simulation results (8).

### Differences in network structure between the 3D and 2D systems

Finally, we investigated the multivalent interactions between AMPAR (or its soluble counterpart, DsRed2. Henceforth, we denote it AMPAR for simplicity), and PSD-95 via TARPs and PDZ domains leads to differences in the network structure between the 3D and 2D systems. As the specific interactions set up throughout this study are treated as virtual association and dissociation reactions, it is possible to specify which TARP specifically interacts with the PDZ domain. Using this property, we examine the distribution of virtual bonds.

We counted the average number of PDZ domains bound per AMPAR from the final 1000 snapshots of the simulation, with 200 of both molecules added to the simulation box. There was a somewhat interesting difference in distribution between the two systems: the results showed that the four TARPs per AMPAR had the highest population in the 3D system, whereas three PDZ domains per AMPAR were found to be the most common in the 2D system (Fig.S5A). This suggests that the protein-domain network in the 3D system exhibits a dense and complicated entangled structure beyond the one-to-one binding of AMPAR complexes and PSD-95s.

To maximize the effect of multivalent interactions, which are the main driving force of phase separation, TARPs were assumed to bind to all four AMPAR subunits in the current work. However, since AMPARs have functional regulatory subunits other than TARPs that competitively bind to the same structural positions as TARPs, this assumption of AMPAR with four TARPs may be unlikely to occur at real synapses (55–57). The results of the virtual bond network analysis imply that even under the condition where the maximum number of TARP (four) is available as an interactive particle, three TARPs are sufficient for AMPARs to form collective structures on the actual postsynaptic membrane, owing to the spatial constraint imposed by the membrane.

The distribution of the number of TARPs bound per PSD-95 also showed slight differences between the two systems: the number of TARPs bound per PSD-95 was almost three in the 3D system, while two were also observed to no small extent in the 2D system (Fig.S5B). Generally, PSD-95 is more readily incorporated into the condensed phase than AMPAR in both systems as there are four TARPs per AMPAR and three PDZ domains per molecule of PSD-95. The setting of the number of molecules may indicate that PSD-95 is relatively more likely to bind more TARPs than the tendency of PDZs to be bound by AMPAR (Fig.S5A), which is consistent with the results for the number of bound TARPs in 3D. However, a surplus of PDZ domains that can interact with TARP was identified in the 2D system despite TARP outnumbering PDZ in the system. Similar to the AMPAR results, this result may be explained by the fact that the number of TARPs that each PSD-95 can access is spatially confined by the geometry of the postsynaptic membrane.

The multiplicity of virtual bonds between a pair of AMPAR and PSD-95 also revealed dimensional differences. The specific interaction between AMPAR and PSD-95 can form up to three virtual bonds (Fig.S5C). The valences of the bonds observed from all pairs of bound AMPAR and PSD-95 pairs were nearly monovalent in both systems, indicating that multi-bonding within the two molecules was relatively rare. However, it was found that the bond valency tended to be higher in the 2D system than in the 3D system (Fig.S5C).

The characteristics of the virtual bond networks in the 3D/2D systems shown in Figs.S4A, B, and C become more prominent when compared with the graphical diagrams (Fig.S5D and S5E).

Figure S4D and S4E illustrate the local bond networks from the final snapshots in the 3D and 2D systems, respectively. These diagrams were drawn starting from AMPAR in the center and tracing 11 bonds (regardless of bond type). For particles shown with a single star, the AMPAR is closest to the center of gravity of the condensed phase in 3D or the largest cluster in 2D. The graphs show that almost all PDZ domains (yellow) are occupied by specific bonds, whereas some TARPs are not used for binding in either system. Notably, the network structure is more developed and densely formed in a 3D system than in a 2D system. The network diagram does not clearly show the difference in the multiplicity of virtual bonds as the formation of multivalent bonds seems to be a relatively rare in both the systems (Fig.S5C). However, the difference in the development of a network is presumed to be correlated with binding multiplicity: in a 3D system, a single molecule, such as AMPAR or PSD-95, can interact with many other molecules. In contrast, the membrane in 2D systems restricts the existing area such that the molecules in and below the membrane can only interact with each other, thus preventing the formation of a dense network. This difference in network structure is hypothesized to be due to the special constraints imposed by the membrane, and it may be a reason why phase separation is less likely to occur in the 2D system than in the 3D system.

## CONCLUSIONS

In this study, to investigate how postsynaptic protein condensates can be formed in and beneath the membrane, we built a mesoscopic model of protein-domain interactions represented by virtual specific reactions and nonspecific interactions and applied it to a mixture of glutamate receptor complexes, AMPAR-TARP, and scaffolding proteins, PSD-95. Based on this model, we could reproduce the phase-separated condensate in the 3D system, which was consistent with the results of the *in vitro* experiment. Molecular simulations revealed a clear distinction between the physical properties of protein assemblies in the 3D and 2D systems. In the 3D system, a soluble variant of the AMPAR-TARP complex and PSD-95 merged into a large complex and produced a liquid-liquid phase-separated condensate. In the 2D system, AMPAR-TARP fixed in the membrane formed clusters with PSD-95, but did not form a stable phase-separated state despite identical domain interactions to the 3D system. Stable phase separation is more difficult to achieve in and under the membrane than in 3D systems.

PSD is a protein condensate formed beneath the postsynaptic membrane and is correlated with signal transmission and synaptic plasticity. This molecular architecture beneath the postsynaptic membrane has been observed using electron microscopy, suggesting that the condensate is stable (10, 16). However, the current simulation results suggest that AMPAR(TARP)_4_ and PSD-95 alone form transient small clusters, but are not sufficient to form a stable phase immediately under the membrane. These are not contradictory as PSD is known to be organized in a hierarchical manner with other scaffolding proteins, such as SAPAP, Shank, and Homer, which are also prone to form protein condensates in *in vitro* 3D systems (8, 58, 59), other than PSD-95, which is generally located in the upper hierarchy of PSD. At the bottom of the PSD hierarchy, the actin cytoskeleton further contributes to retaining the protein network and anchors the condensate to the cell body. With this hierarchy, protein networks may physically be between 2D and 3D systems, which could enhance the stability of the condensate. Incorporation of these additional scaffolding and cytoskeleton proteins into mesoscopic computational modeling would be future challenge.

More generally, we suggest that qualitative aspect of the current results is applicable to other protein assemblies with multidomain interactions, since the mesoscale model we employed only depends on a limited number of parameters and is less protein-specific. Further research may explore a more general consideration of the extent to which domain multi-valency and protein-protein affinity affect phase separation.

In general, the phase transition depends on physical dimensions (9, 60, 61). One-dimensional systems with finite-range interactions cannot usually show phase transition. 3D systems often exhibit sharp phase transitions. 2D systems are marginal. For example, the Ising model shows the phase transitions in 2D and 3D systems, whereas the Heisenberg model of ferromagnetism shows the phase transitions only in 3D, but not in 2D systems. The current results are in harmony with the general theories of phase transitions. In recent studies, LLPS has often been observed in *in vitro* 3D systems. It should be noted that the observation of spherical droplets in these 3D systems does not necessarily indicate the stable phase separation of systems that are restricted to 2D.

## AUTHOR CONTRIBUTIONS

RY and ST conceived and designed the study. RY constructed a mesoscopic model of proteins in PSD, set up the simulation program ReaDDy 2, performed the molecular simulation, and conducted the analysis. RY and ST wrote the manuscript.

### DECLARATION OF INTERESTS

The authors declare no competing interests.

ACKNOWLEDGEMENTS

We would like to thank Yutaka Murata, Giovanni Brandani, and Yasunori Hayashi for the insightful discussions. This work was supported partly by JSPS KAKENHI grants 20H05934 (ST) and 21H02441 (ST) and by the Japan Science and Technology Agency (JST) grant JPMJCR1762 (ST).

## SUPPORTING MATERIAL

Figure S1. Convergence examination.

Figure S2. Preparation of the initial configuration in the reverse simulation in Fig.6 and the coefficients of variation in two systems.

Figure S3. Simulation in a 2D system with a slab box.

Figure S4. Size scaling analysis with planar geometry. Figure S5. Network analysis in 3D and 2D systems.

Movie S1. Phase separation dynamics for a mixture of 200 PSD-95 and DsRed2(TARPc)_4_ complexes in a 3D system.

Movie S2. Molecular assembly dynamics for a mixture of 50 palmitoylated PSD-95 and AMPAR(TARP)_4_ in and under the membrane.

## REFERENCES

1. Wang, B., L. Zhang, T. Dai, Z. Qin, H. Lu, L. Zhang, and F. Zhou. 2021. Liquid-liquid phase separation in human health and diseases. Signal Transduct Target Ther. 6:290.

2. Wu, H., and M. Fuxreiter. 2016. The Structure and Dynamics of Higher-Order Assemblies: Amyloids, Signalosomes, and Granules. Cell. 165:1055–1066.

3. Das, S., Y.-H. Lin, R.M. Vernon, J.D. Forman-Kay, and H.S. Chan. 2020. Comparative roles of charge, π, and hydrophobic interactions in sequence-dependent phase separation of intrinsically disordered proteins. Proc Natl Acad Sci U S A. 117:28795–28805.

4. Murthy, A.C., G.L. Dignon, Y. Kan, G.H. Zerze, S.H. Parekh, J. Mittal, and N.L. Fawzi. 2019. Molecular interactions underlying liquid−liquid phase separation of the FUS low-complexity domain. Nat Struct Mol Biol. 26:637–648.

5. Dignon, G.L., U. States, and R.B. Best. 2021. Biomolecular Phase Separation: From Molecular Driving Forces to Macroscopic Properties. Annu Rev Phys Chem. 71:53–75.

6. Brangwynne, C.P., C.R. Eckmann, D.S. Courson, A. Rybarska, C. Hoege, J. Gharakhani, F. Jülicher, and A.A. Hyman. 2009. Germline P granules are liquid droplets that localize by controlled dissolution/condensation. Science. 324:1729–32.

7. Li, P., S. Banjade, H.-C. Cheng, S. Kim, B. Chen, L. Guo, M. Llaguno, J. v Hollingsworth, D.S. King, S.F. Banani, P.S. Russo, Q.-X. Jiang, B.T. Nixon, and M.K. Rosen. 2012. Phase transitions in the assembly of multivalent signalling proteins. Nature. 483:336–40.

8. Zeng, M., X. Chen, D. Guan, J. Xu, H. Wu, P. Tong, and M. Zhang. 2018. Reconstituted Postsynaptic Density as a Molecular Platform for Understanding Synapse Formation and Plasticity. Cell. 174:1172–1187.e16.

9. Huang, K. 1987. Statistical Mechanics. 2nd Edition. John Wiley & Sons, Inc.

10. Palay, S.L. 1956. Synapses in the central nervous system. J Biophys Biochem Cytol. 2:193–202.

11. Harris, K.M., F.E. Jensen, and B. Tsao. 1992. Three-dimensional structure of dendritic spines and synapses in rat hippocampus (CA1) at postnatal day 15 and adult ages: implications for the maturation of synaptic physiology and long-term potentiation. J Neurosci. 12:2685–705.

12. Husi, H., M.A. Ward, J.S. Choudhary, W.P. Blackstock, and S.G. Grant. 2000. Proteomic analysis of NMDA receptor-adhesion protein signaling complexes. Nat Neurosci. 3:661–9.

13. Peng, J., M.J. Kim, D. Cheng, D.M. Duong, S.P. Gygi, and M. Sheng. 2004. Semiquantitative proteomic analysis of rat forebrain postsynaptic density fractions by mass spectrometry. J Biol Chem. 279:21003–11.

14. Yoshimura, Y., Y. Yamauchi, T. Shinkawa, M. Taoka, H. Donai, N. Takahashi, T. Isobe, and T. Yamauchi. 2004. Molecular constituents of the postsynaptic density fraction revealed by proteomic analysis using multidimensional liquid chromatography-tandem mass spectrometry. J Neurochem. 88:759–68.

15. Cheng, D., C.C. Hoogenraad, J. Rush, E. Ramm, M.A. Schlager, D.M. Duong, P. Xu, S.R. Wijayawardana, J. Hanfelt, T. Nakagawa, M. Sheng, and J. Peng. 2006. Relative and absolute quantification of postsynaptic density proteome isolated from rat forebrain and cerebellum. Mol Cell Proteomics. 5:1158–70.

16. Carlin, R.K., D.J. Grab, R.S. Cohen, and P. Siekevitz. 1980. Isolation and characterization of postsynaptic densities from various brain regions: enrichment of different types of postsynaptic densities. J Cell Biol. 86:831–45.

17. Collins, M.O., L. Yu, M.P. Coba, H. Husi, I. Campuzano, W.P. Blackstock, J.S. Choudhary, and S.G.N. Grant. 2005. Proteomic analysis of in vivo phosphorylated synaptic proteins. J Biol Chem. 280:5972–82.

18. Siekevitz, P. 1985. The postsynaptic density: a possible role in long-lasting effects in the central nervous system. Proc Natl Acad Sci U S A. 82:3494–8.

19. Chen, X., L. Vinade, R.D. Leapman, J.D. Petersen, T. Nakagawa, T.M. Phillips, M. Sheng, and T.S. Reese. 2005. Mass of the postsynaptic density and enumeration of three key molecules. Proc Natl Acad Sci U S A. 102:11551–11556.

20. Sugiyama, Y., I. Kawabata, K. Sobue, and S. Okabe. 2005. Determination of absolute protein numbers in single synapses by a GFP-based calibration technique. Nat Methods. 2:677–684.

21. Zhang, M., and W. Wang. 2003. Organization of signaling complexes by PDZ-domain scaffold proteins. Acc Chem Res. 36:530–8.

22. Liu, P.W., T. Hosokawa, and Y. Hayashi. 2021. Regulation of synaptic nanodomain by liquid–liquid phase separation: A novel mechanism of synaptic plasticity. Curr Opin Neurobiol. 69:84–92.

23. Glavinovíc, M.I., and H.R. Rabie. 1998. Monte Carlo simulation of spontaneous miniature excitatory postsynaptic currents in rat hippocampal synapse in the presence and absence of desensitization. Pflugers Arch. 435:193–202.

24. Franks, K.M., T.M. Bartol, and T.J. Sejnowski. 2002. A Monte Carlo model reveals independent signaling at central glutamatergic synapses. Biophys J. 83:2333–48.

25. Raghavachari, S., and J.E. Lisman. 2004. Properties of quantal transmission at CA1 synapses. J Neurophysiol. 92:2456–67.

26. Heine, M., L. Groc, R. Frischknecht, J.-C. Béïque, B. Lounis, G. Rumbaugh, R.L. Huganir, L. Cognet, and D. Choquet. 2008. Surface mobility of postsynaptic AMPARs tunes synaptic transmission. Science. 320:201–5.

27. Nair, D., E. Hosy, J.D. Petersen, A. Constals, G. Giannone, D. Choquet, and J.-B. Sibarita. 2013. Super-resolution imaging reveals that AMPA receptors inside synapses are dynamically organized in nanodomains regulated by PSD95. J Neurosci. 33:13204–24.

28. Zeng, M., J. Díaz-Alonso, F. Ye, X. Chen, J. Xu, Z. Ji, R.A. Nicoll, and M. Zhang. 2019. Phase Separation-Mediated TARP/MAGUK Complex Condensation and AMPA Receptor Synaptic Transmission. Neuron. 104:529–543.e6.

29. Hosokawa, T., P.-W. Liu, Q. Cai, J.S. Ferreira, F. Levet, C. Butler, J.-B. Sibarita, D. Choquet, L. Groc, E. Hosy, M. Zhang, and Y. Hayashi. 2021. CaMKII activation persistently segregates postsynaptic proteins via liquid phase separation. Nat Neurosci. 24:777–785.

30. Chattaraj, A., M. Youngstrom, and L.M. Loew. 2019. The Interplay of Structural and Cellular Biophysics Controls Clustering of Multivalent Molecules. Biophys J. 116:560–572.

31. Chattaraj, A., M.L. Blinov, and L.M. Loew. 2021. The solubility product extends the buffering concept to heterotypic biomolecular condensates. Elife. 10:e67176.

32. Hoffmann, M., C. Fröhner, and F. Noé. 2019. ReaDDy 2: Fast and flexible software framework for interacting-particle reaction dynamics. PLoS Comput Biol. 15:e1006830.

33. Su, Z., K. Dhusia, and Y. Wu. 2020. Understand the Functions of Scaffold Proteins in Cell Signaling by a Mesoscopic Simulation Method. Biophys J. 119:2116–2126.

34. Chen, L., D.M. Chetkovich, R.S. Petralia, N.T. Sweeney, Y. Kawasaki, R.J. Wenthold, D.S. Bredt, and R.A. Nicoll. 2000. Stargazin regulates synaptic targeting of AMPA receptors by two distinct mechanisms. Nature. 408:936–43.

35. Hafner, A.-S., A.C. Penn, D. Grillo-Bosch, N. Retailleau, C. Poujol, A. Philippat, F. Coussen, M. Sainlos, P. Opazo, and D. Choquet. 2015. Lengthening of the Stargazin Cytoplasmic Tail Increases Synaptic Transmission by Promoting Interaction to Deeper Domains of PSD-95. Neuron. 86:475–89.

36. Topinka, J.R., and D.S. Bredt. 1998. N-terminal palmitoylation of PSD-95 regulates association with cell membranes and interaction with K+ channel Kv1.4. Neuron. 20:125–34.

37. Dill, K.A., K. Ghosh, and J.D. Schmit. 2011. Physical limits of cells and proteomes. Proc Natl Acad Sci U S A. 108:17876–82.

38. Saito, H., T. Araiso, H. Shirahama, and T. Koyama. 1991. Dynamics of the Bilayer-Water Interface of Phospholipid Vesicles and the Effect of Cholesterol: A Picosecond Fluorescence Anisotropy Study. The Journal of Biochemistry. 109:559–565.

39. Smoluchowski, M. v. 1916. Drei Vortrage uber Diffusion, Brownsche Bewegung und Koagulation von Kolloidteilchen. Zeitschrift fur Physik. 17:557–585.

40. Schöneberg, J., and F. Noé. 2013. ReaDDy--a software for particle-based reaction-diffusion dynamics in crowded cellular environments. PLoS One. 8:e74261.

41. Vistrup-Parry, M., X. Chen, T.L. Johansen, S. Bach, S.C. Buch-Larsen, C.R.O. Bartling, C. Ma, L.S. Clemmensen, M.L. Nielsen, M. Zhang, and K. Strømgaard. 2021. Site-specific phosphorylation of PSD-95 dynamically regulates the postsynaptic density as observed by phase separation. iScience. 24:103268.

42. Dignon, G.L., W. Zheng, R.B. Best, Y.C. Kim, and J. Mittal. 2018. Relation between single-molecule properties and phase behavior of intrinsically disordered proteins. Proc Natl Acad Sci U S A. 115:9929–9934.

43. Jung, H., and A. Yethiraj. 2018. A simulation method for the phase diagram of complex fluid mixtures. J Chem Phys. 148:244903.

44. McSwiggen, D.T., M. Mir, X. Darzacq, and R. Tjian. 2019. Evaluating phase separation in live cells: diagnosis, caveats, and functional consequences. Genes Dev. 33:1619–1634.

45. Murata, Y., T. Niina, and S. Takada. 2022. The stoichiometric interaction model for mesoscopic MD simulations of liquid-liquid phase separation. Biophys J. 121:4382–4393.

46. Lin, Y.-H., H. Wu, B. Jia, M. Zhang, and H.S. Chan. 2022. Assembly of model postsynaptic densities involves interactions auxiliary to stoichiometric binding. Biophys J. 121:157–171.

47. Axelrod, D., D.E. Koppel, J. Schlessinger, E. Elson, and W.W. Webb. 1976. Mobility measurement by analysis of fluorescence photobleaching recovery kinetics. Biophys J. 16:1055–69.

48. Meyvis, T.K., S.C. de Smedt, P. van Oostveldt, and J. Demeester. 1999. Fluorescence recovery after photobleaching: a versatile tool for mobility and interaction measurements in pharmaceutical research. Pharm Res. 16:1153–62.

49. Taylor, N.O., M.-T. Wei, H.A. Stone, and C.P. Brangwynne. 2019. Quantifying Dynamics in Phase-Separated Condensates Using Fluorescence Recovery after Photobleaching. Biophys J. 117:1285–1300.

50. Nusser, Z., R. Lujan, G. Laube, J.D. Roberts, E. Molnar, and P. Somogyi. 1998. Cell type and pathway dependence of synaptic AMPA receptor number and variability in the hippocampus. Neuron. 21:545–59.

51. Tanaka, J., M. Matsuzaki, E. Tarusawa, A. Momiyama, E. Molnar, H. Kasai, and R. Shigemoto. 2005. Number and density of AMPA receptors in single synapses in immature cerebellum. J Neurosci. 25:799–807.

52. Masugi-Tokita, M., E. Tarusawa, M. Watanabe, E. Molnár, K. Fujimoto, and R. Shigemoto. 2007. Number and density of AMPA receptors in individual synapses in the rat cerebellum as revealed by SDS-digested freeze-fracture replica labeling. J Neurosci. 27:2135–44.

53. Onuki, A. 2002. Phase Transition Dynamics. Cambridge University Press.

54. Tateno, M., and H. Tanaka. 2021. Power-law coarsening in network-forming phase separation governed by mechanical relaxation. Nat Commun. 12:912.

55. Kamalova, A., and T. Nakagawa. 2021. AMPA receptor structure and auxiliary subunits. J Physiol. 599:453–469.

56. Shi, Y., W. Lu, A.D. Milstein, and R.A. Nicoll. 2009. The stoichiometry of AMPA receptors and TARPs varies by neuronal cell type. Neuron. 62:633–40.

57. Schwenk, J., N. Harmel, A. Brechet, G. Zolles, H. Berkefeld,C.S. Müller, W. Bildl, D. Baehrens, B. Hüber, A. Kulik, N. Klöcker, U. Schulte, and B. Fakler. 2012. High-resolution proteomics unravel architecture and molecular diversity of native AMPA receptor complexes. Neuron. 74:621–33.

58. Sheng, M., and C.C. Hoogenraad. 2007. The postsynaptic architecture of excitatory synapses: a more quantitative view. Annu Rev Biochem. 76:823–47.

59. Feng, Z., X. Wu, and M. Zhang. 2021. Presynaptic bouton compartmentalization and postsynaptic density-mediated glutamate receptor clustering via phase separation. Neuropharmacology. 193:108622.

60. Gallavotti, G. 1972. The phase separation line in the two-dimensional Ising model. Commun Math Phys. 27:103–136.

61. Emery, V., S. Kivelson, and H. Lin. 1990. Phase separation in the t-J model. Phys Rev Lett. 64:475–478.

