## Supplemental information for "Postsynaptic protein assembly in three- and two-dimensions studied by mesoscopic simulations"

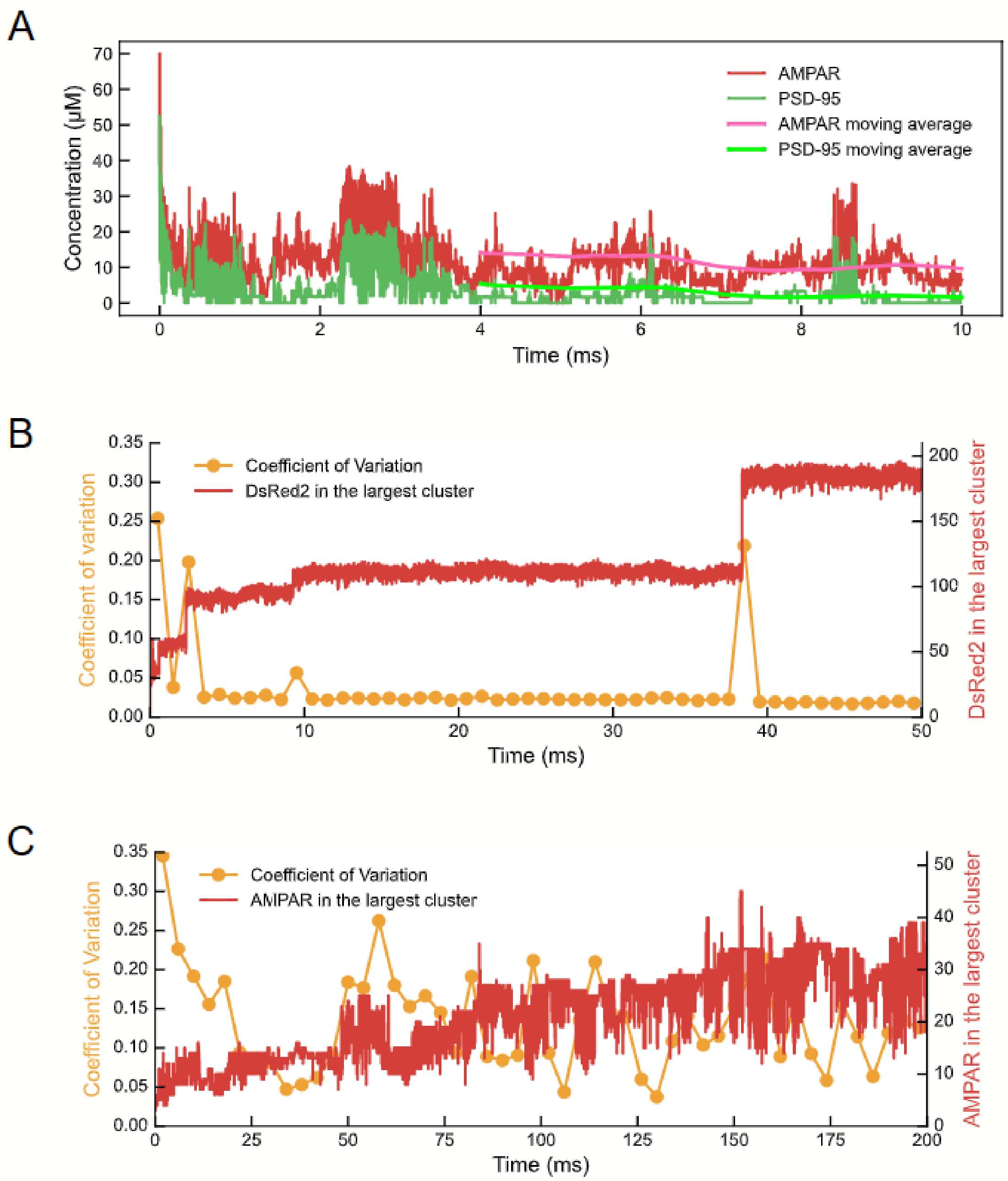


**Figure S1. Convergence examination.**

A) Regarding the convergence of simulations in Fig.2B, the time courses of the concentrations of AMPAR (red) and PSD-95 (green) in the dilute phase were shown for the case of $(E_{\mathrm{nonspe}}, G_{spe}) = (28.7, 1.5)$ (kJ/mol). The simulation started from the completely phase-separated state. In addition to raw time course, the moving average over 4 ms were plotted (the average from t-4 ms to t ms was plotted at t). The results suggest the convergence up to 10 ms. B) Regarding the trajectory in Fig.3AB, time series of the number of DsRed2s in the largest cluster (red) and the coefficient of variation in every 1 ms (orange) in the 3D system. C) Time series of the number of AMPARs in the largest cluster (red) and the coefficient of variation in every 4 ms (orange) in 2D system. Both of the time in these figures are plotted on linear scale.

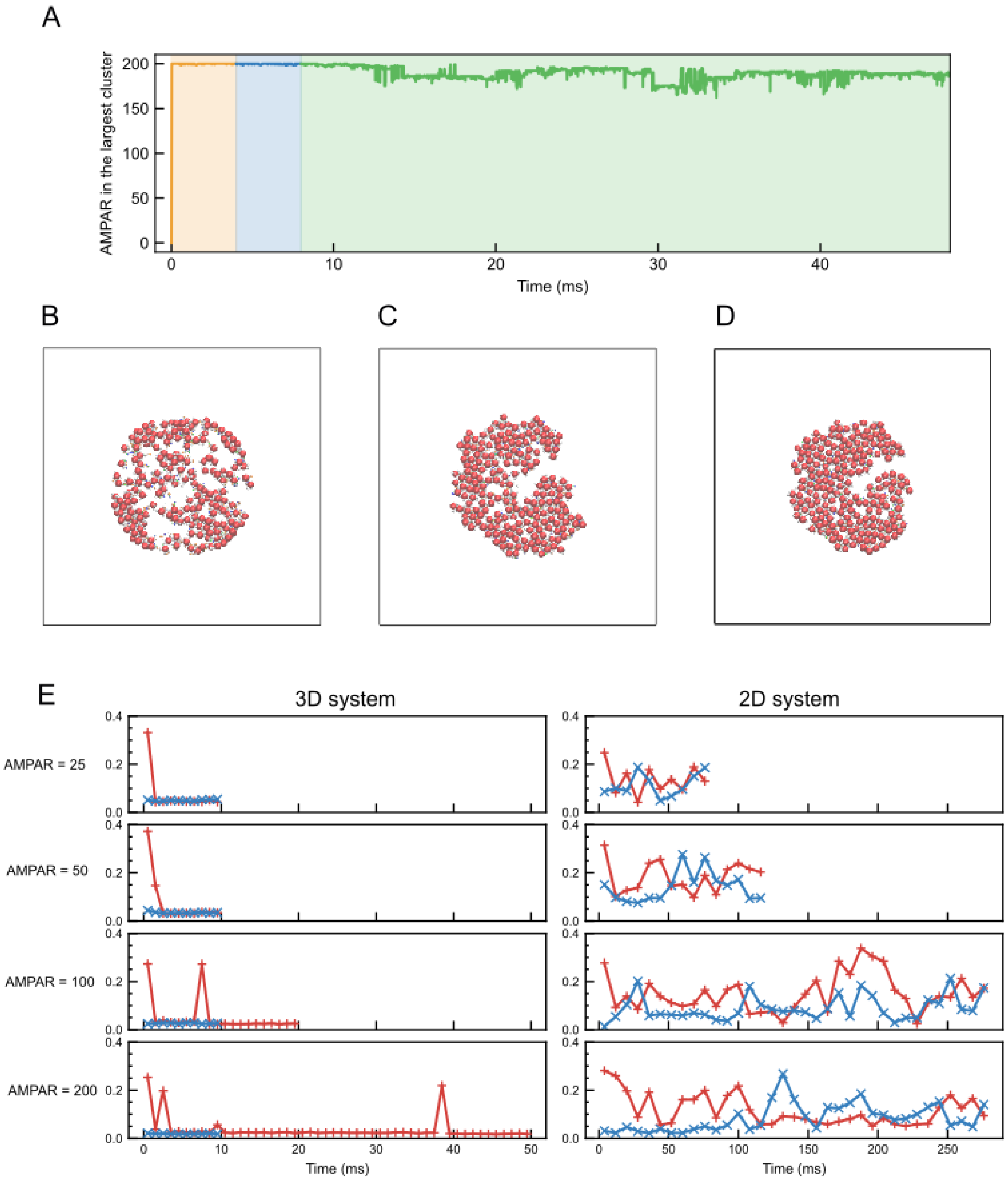
A
AMPAR in the largest cluster
Time (ms)
B
C
D
E
3D system
2D system
AMPAR = 25
AMPAR = 50
AMPAR = 100
AMPAR = 200
Time (ms)
Time (ms)

**Figure S2. Preparation of the initial configuration in the reverse simulation in Fig. 6 and the coefficients of variation in two systems**

A) To create an initial configuration for the reverse simulation, we employed a two-step procedure. First, we ran a preliminary simulation for $4 \times 10^6$ MD steps (propagation step, orange line), in which all the molecules in and under the membrane were first restrained to a circle of radius $7.5\sqrt{n_{AMPAR}}\,nm$ in the xy plane. $n_{AMPAR}$ is the number of molecules in a system. During this simulation, we applied a virtual association reaction for a specific interaction but not the dissociation reaction. Second, we removed the circular restraint and performed another $4 \times 10^6$ MD step simulation (relaxation step, blue line) with an association reaction alone to relax the configuration. Finally, we initiated the reverse simulation applying both association and dissociation reactions (reverse step, green line). Propagation and relaxation step are not included in trajectories shown in the main figure. B) Snapshot of the initial structure (t = 0 ms). C) Snapshot after the preliminary simulation (t = 4 ms). D) Snapshot after the relaxation step (t = 8 ms). Starting with this configuration, we performed the reverse simulation as described in Methods. E) Comparison of coefficient of variation between 3D and 2D system. The crosses (red) indicate the results of the forward simulation and the x-mark (blue) indicate the results of the reverse simulation. Means and variances were measured and calculated every 1 ms in 3D system and every 8 ms in 2D system.

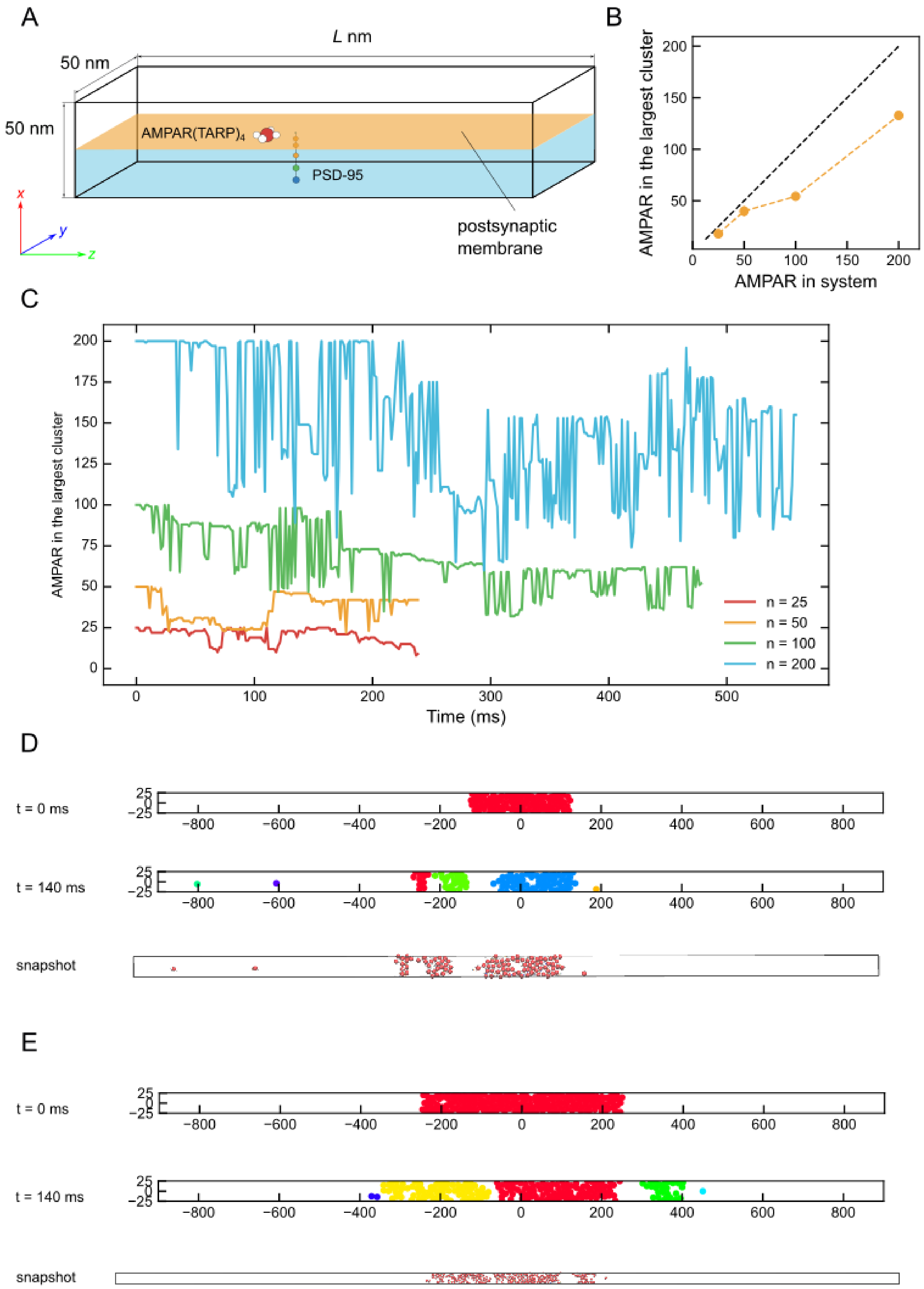
A
L nm
50 nm
50 nm
AMPAR(TARP)4
PSD-95
postsynaptic membrane
x
y
z
B
AMPAR in the largest cluster
AMPAR in system
C
AMPAR in the largest cluster
Time (ms)
n = 25
n = 50
n = 100
n = 200
D
t = 0 ms
t = 140 ms
snapshot
E
t = 0 ms
t = 140 ms
snapshot

**Figure S3. Simulation in a 2D system with a slab box**

To assess the finite-size effect of the simulation box, we also performed a 2D simulation in a 2D-variant of the slab box. A) The 2D slab box. Particles representing AMPAR, TARP, and N-terminal of PSD-95 diffuse on the planar membrane (orange), while other particles diffuse in the cytosol just below the membrane (sky-blue). The length of z-axis of slab box ($L$) is determined to be the same as the areal density at the dense condition in the 2D simulation. Specifically, $L$ is 1800 nm and 3600 nm when the number of AMPARs in the system is 100 and 200, respectively. B) The mean of the largest AMPAR cluster size as a function on the total number of AMPARs in the 2D system with a slab box. Black dashed line is the line assumed that the cluster growth is proportional to the cluster size in the system in Fig.6C. C) The reverse trajectories of the largest AMPAR cluster size as a function of time for two different system sizes of self-similarity; 25(red), 50(orange), 100(green) and 200(blue) AMPAR(TARP)$_4$s and PSD-95s with the dense condition. D) Cluster structures at t = 0 ms (top), at t = 140 ms (middle), and a snapshot at t = 140 ms (bottom) with 100 AMPAR(TARP)$_4$s and 100 PSD-95s in the system. E) Cluster structure at t = 0 ms (top), at t = 140 ms (middle), and snapshot at t = 140 ms (bottom) with 200 AMPAR(TARP)$_4$s and 200 PSD-95s in the system. In the cluster structure in Fig.S2D and E, AMPARs belonging to the same cluster are classified and plotted in the same color.

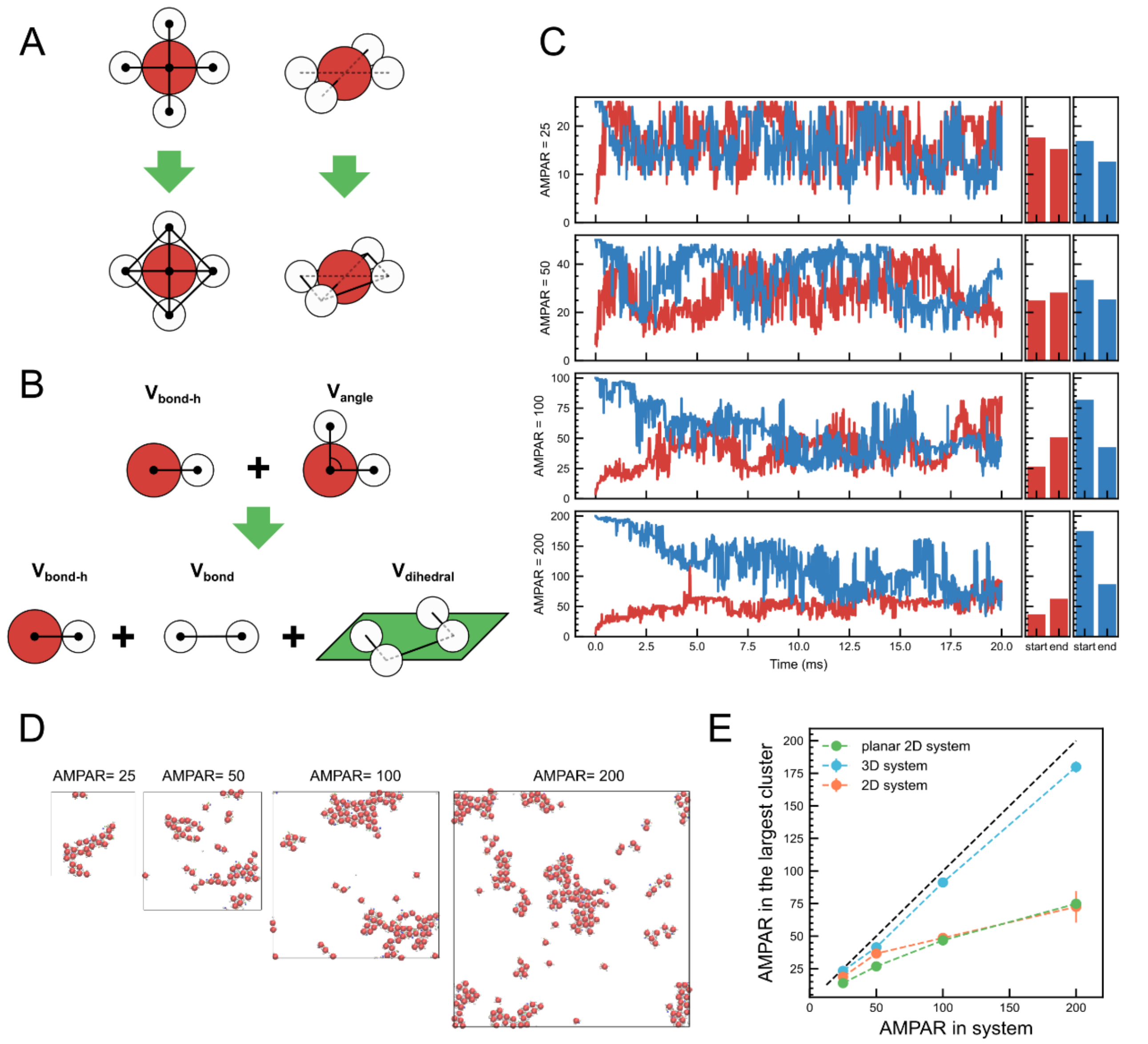


**Figure S4. Size scaling analysis with planar geometry**

Size scaling analysis in 2D system with planar TARP geometry. A) AMPAR(TARP)$_4$ topologic bonds applied in 2D system (top) and planar 2D system (bottom). B) AMPAR(TARP)$_4$ potentials applied in 2D system (top) and planar 2D system (bottom). C) The forward (red) and reverse (blue) trajectories of the largest AMPAR cluster size as a function of time for four different system sizes of self-similarity; 25, 50, 100, and 200 AMPAR(TARP)$_4$s and PSD-95s for the 2D systems (left panel). To allow the protein assembly to reach equilibrium quickly, the viscosity of the membrane was set to 20 times that used in the 2D system in Figure 6. This modification is expected to have little effect on the final number of molecules in the largest cluster since the diffusion coefficient theoretically does not affect the equilibrium state of the molecular assembly. The mean of the largest clusters at the beginning (denoted as start) and the end (denoted as end) of each simulation (right panel). D) The final snapshots of the four systems. E) The mean of the largest AMPAR cluster size as a function on the total number of AMPARs with planar geometry (green). Black dashed line is the line assumed that the cluster growth is proportional to the cluster size in the system.

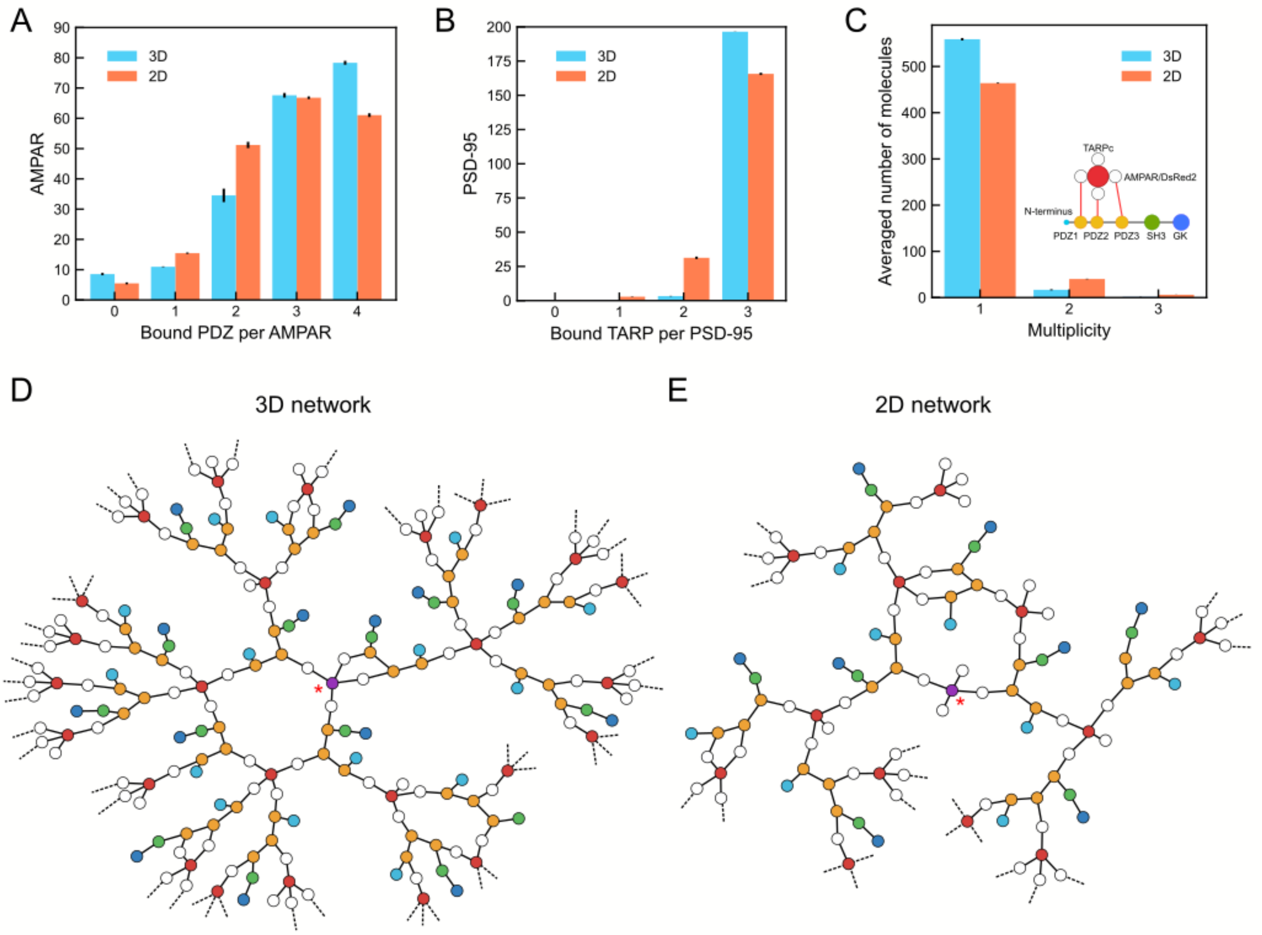


**Figure S5. Network analysis in 3D and 2D systems**

The interaction networks in the 3D and 2D systems. A) The average number of PDZ domains bound per DsRed2 in 3D (blue) and AMPAR in 2D (coral) systems. B) The average number of TARPc's and TARPs bound per PSD-95 in the 3D (blue) and 2D (coral) systems, respectively. C) The multiplicity of the virtual bond between each pair of an DsRed2 (AMPAR) and a PSD-95 in the 3D (blue) and 2D (coral) systems. D) Representative local network structures near the center (left) and in periphery (right) of the 3D condensed phase. E) Representative local network structures near the center (left) and in periphery (right) of the 2D cluster. In D and E, purple molecules with single star are those closest to the geometric center of the condensed phase in 3D system and largest cluster in 2D system, respectively. Data for the 2D system are from the case of 200 AMPAR(TARP)$_4$'s in the dense condition. Dashed line denotes that there is a further network structure continuing from the molecule.
